# Multicomponent nature underlies the extraordinary mechanical properties of spider dragline silk

**DOI:** 10.1101/2021.04.22.441049

**Authors:** Nobuaki Kono, Hiroyuki Nakamura, Masaru Mori, Yuki Yoshida, Rintaro Ohtoshi, Ali D Malay, Daniel A Pedrazzoli Moran, Masaru Tomita, Keiji Numata, Kazuharu Arakawa

## Abstract

Dragline silk of golden orb-weaver spiders (Nephilinae) is noted for its unsurpassed toughness, combining extraordinary extensibility and tensile strength, suggesting industrial application as a sustainable biopolymer material. To pinpoint the molecular composition of dragline silk and the roles of its constituents in achieving its mechanical properties, we report a multiomics approach combining high-quality genome sequencing and assembly, silk gland transcriptomics, and dragline silk proteomics of four Nephilinae spiders. We observed the consistent presence of the MaSp3B spidroin unique to this subfamily, as well as several non-spidroin SpiCE proteins. Artificial synthesis and combination of these components in vitro showed that the multicomponent nature of dragline silk, including MaSp3B and SpiCE, along with MaSp1 and MaSp2, is essential to realize the mechanical properties of spider dragline silk.

## Introduction

Spider silk is a typical natural high-performance structural protein and has potential for numerous applications as a protein biopolymer with biodegradability and biocompatibility (1). Spider silk has a tensile strength superior to that of steel, yet it is highly elastic, showing greater toughness than aramid fibers such as Kevlar (2); thus, it has received interest for use in industrial applications (1, 3). Orb-weaver spiders, especially those belonging to the family Araneidae, are often used as models in natural spider silk research. The average mechanical properties of their dragline silks reach approximately 1 GPa for breaking strength, 30% for breaking strain, and 130 MJ/m^3^ for toughness (4). Numerous works have reported the use of recombinant protein and artificial fiber spinning to produce artificial spider silk in genetically optimized organisms (5–10), but it remains challenging to fully reproduce and rival the mechanical properties of natural silks (6, 11–13). The difficulties are multifactorial; the unusually large protein size and the repetitive nature of its sequence are inherent challenges for synthesis, and the natural conditions of spinning are only beginning to be fully uncovered (13–15). However, one primary reason that full reproduction has not been successful is that the previous recombinant approaches employed only MaSp1, only MaSp2, or at most a combination of the two (11, 16–18). In fact, it is often suggested that dragline silk is composed primarily of two components, MaSp1 and MaSp2 (5, 19–23). However, recent proteome analyses suggest the existence of additional components in spider silks, such as a cysteine-rich protein (CRP) in black widows (24–26). Genome and transcriptome analyses have identified many MaSp families (27, 28), and in the genus *Araneus*, it is known that dragline silk contains nearly equal amounts of MaSp3 and MaSp1/2 (28). Furthermore, low molecular weight (LMW) non-spidroin proteins, such as spider-silk constituting element (SpiCE), have been found by transcriptomic and proteomic analyses. SpiCE is a protein of unknown function that is commonly highly expressed in the silk gland and in spider silk. It is becoming apparent that dragline silk is a complex multicomponent material containing much more than MaSp1 and MaSp2.

To pinpoint the protein constituents of dragline silk through quantitative proteomics, a high-quality reference genome and full-length coding sequence annotation are essential to allow the correct and comprehensive identification of the proteins corresponding to the detected peptide fragments. Since spider fibroin genes are extremely long (∼10 kbp) and consist almost entirely of repeat sequences (29, 30), genome sequencing using PCR-free long reads is critical, and in order to eliminate false-positive annotations, predicted coding sequences need to be confirmed on the bases of conservation in closely related species and actual mRNA expression in the silk gland. Hence, we took a multiomics approach to quantitatively identify the dragline protein constituents and the genomes of four golden silk orb-weavers (subfamily Nephilinae): *Trichonephila clavata*, *Trichonephila clavipes*, *Trichonephila inaurata madagascariensis*, and *Nephila pilipes.* These spiders are reported to produce high-performance dragline silk with average toughness values of 169, 131, 285, and 292 MJ/m^3^, respectively (Fig. S1). *T. clavipes* genome data have already been reported (27), but we chose to construct *ab initio* assemblies, including for this species, since the existing genome is based on PCR-amplified sequencing for fibroins, and some fibroin gene sequences remain incomplete. Moreover, the existing *T. clavipes* assembly is suspected to be substantially contaminated, as the entirety of the longest scaffold has been identified as bacterial in origin (31).

## Results

### Draft genomes of four Nephilinae spiders

Spider genomes are large and complex, and de novo sequencing is challenging. We extracted genomic DNA from dissected leg and cephalothorax tissue of adult female individuals, conducted hybrid sequencing with short- and long-read sequence technologies (32) (see Methods). The 10X Genomics linked sequencing produced 702.57 M reads (106.09 Gb), 1,090 M reads (164.60 Gb), 1,010.42 M reads (152.57 Gb), and 900.42 M reads (135.96 Gb) from *T. clavata*, *T. clavipes*, *T. inaurata madagascariensis*, and *N. pilipes*, respectively (S1 Table). PacBio sequencing for the *T. clavata* genome yielded 74.81 Gb of data with an average length of 8.24 kb and an N50 of 11.1 kb (S1 Table). Furthermore, 2.19 M reads (11.03 Gb), 4.70 M reads (41.18 Gb), 5.88 M reads (24.33 Gb), 2.92 M reads (8.48 Gb) were generated by Nanopore sequencing of gDNA from *T. clavata*, *T. clavipes*, *T. inaurata madagascariensis*, and *N. pilipes*, respectively (S1 Table). Total coverage was approximately 80X. Obtained sequence reads were assembled, polished, corrected, and contaminant eliminated (see Methods) the resulting the number of scaffolds and N50 length in the *T. clavata*, *T. clavipes*, *T. inaurata madagascariensis*, and *N. pilipes* draft genomes were 39,792 (N50 of 112,957 bp), 21,438 (N50 of 738,925 bp), 28,263 (N50 of 231,352 bp), and 132,899 (N50 of 292,147 bp), respectively (Table 1, Fig. S3). The completeness of each assembled result was assessed by BUSCO (33). The completeness scores were 90.6%, 93.0%, 89.8%, and 92.9% for the eukaryote lineage (Table 1), indicating the high quality of the obtained draft genomes. The tRNAs and ribosomal RNAs were annotated using tRNAscan-SE v2.0 (34) and Barrnap (https://github.com/tseemann/barrnap) with default parameters. In total, 8,008, 11,433, 8,679, and 1,837 tRNAs, and 248 (49 18S, 49 28S, 43 5.8S, and 107 5S), 106 (15 18S, 5 28S, 9 5.8S, and 77 5S), 128 (11 18S, 10 28S, 5 5.8S, and 102 5S), and 167 (8 18S, 8 28S, 6 5.8S, and 29 5S) rRNA copies were predicted in the *T. clavata*, *T. clavipes*, *T. inaurata madagascariensis*, and *N. pilipes* genomes, respectively. The *T. clavipes* genome improves upon the previous assembly (27) in quality and completeness (longest contig, 4.65 Mbp; N50 length, 738 kbp) (Fig. S4 and SI Text).

**Table 1.**
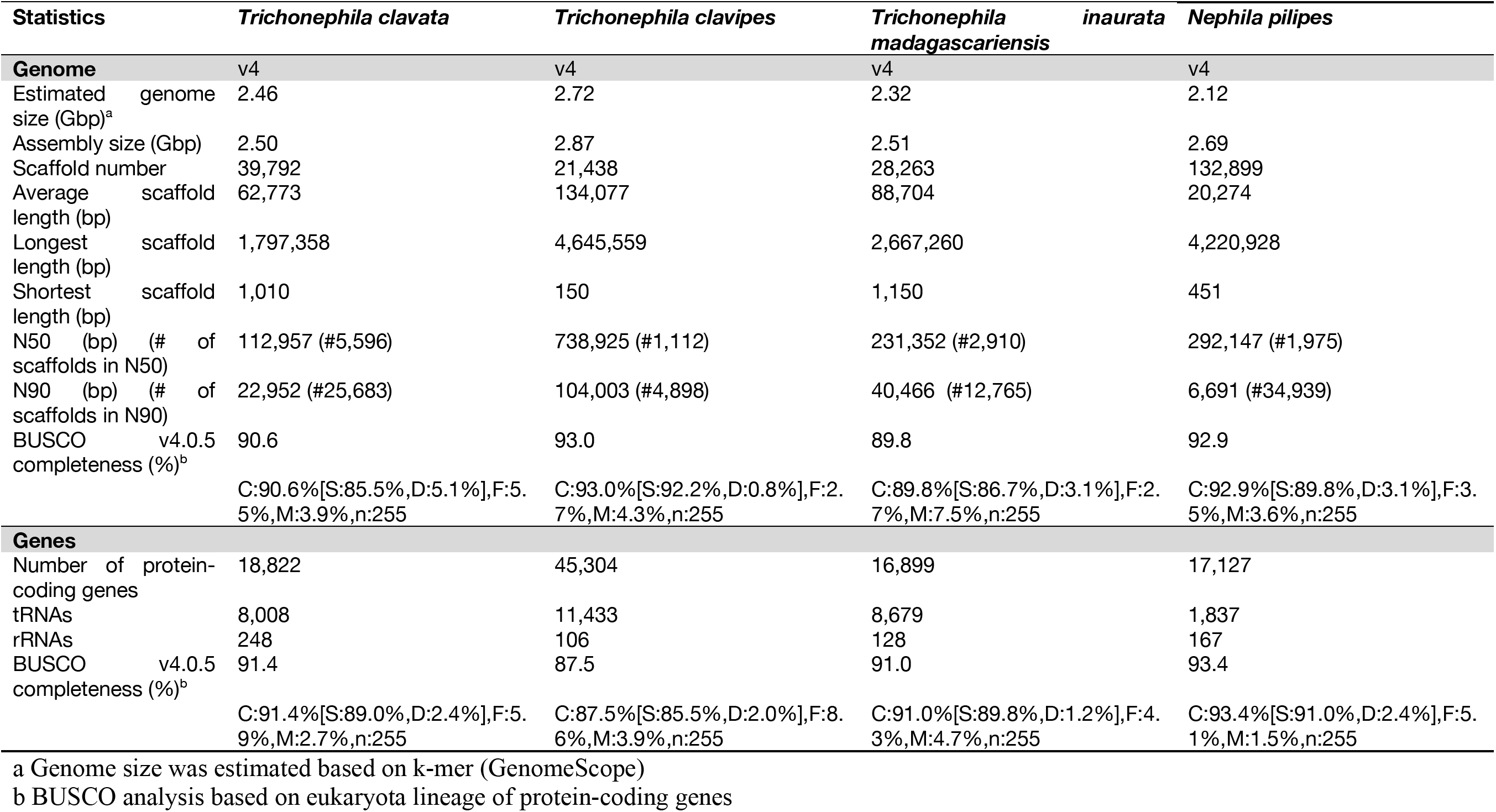
Assembly and annotation statistics of four spider draft genomes (T. clavata, T. clavipes, T. inaurata madagascariensis, and N. pilipes).

The number of protein-coding genes predicted by BRAKER was initially 70,418, 517,373, 51,581, and 72,271. Redundant genes were eliminated by CD-HIT-EST (35) clustering with a nucleotide identity of 97%. To obtain a functional gene set, we removed the genes with an expression level of less than 0.1 and unannotated genes. Finally, 18,822, 45,304, 16,899, and 17,127 functional protein-coding gene sets were obtained for *T. clavata*, *T. clavipes*, *T. inaurata madagascariensis*, and *N. pilipes,* respectively (Table 1). BUSCO (v4.0.5) was used to determine the quality of our functional gene set using the eukaryote lineage, and the completeness scores were 91.4%, 87.5%, 91.0%, and 93.4% for the eukaryote lineage (Table 1).

### Catalog of spidroins in four Nephilinae spiders

Spidroin genes are structured with a long repeat domain sandwiched between the nonrepetitive N/C-terminal domains and an extreme length, of 10 kbp or more (29, 30, 36, 37). Short-read sequencing and PCR amplification technology alone often cause fragmentation, collapse, and chimerization of spidroin genes. Therefore, long-read sequencing using MinION or GridION is an essential technique for finding spidroins because these technologies can sequence a region covering the entire length of the spidroin-coding gene. However, the sequence of the repeat region is very delicate, and the sequence quality of long reads alone is not sufficiently high. We have contributed to the construction of the spidroin catalog using hybrid sequencing, which combines various sequencing technologies and has been reported previously (28, 32, 38). In a previous study, we reported the spidroin catalog, including complete spidroin CDS in *A. ventricosus*, and showed that known spidroin genes that were sequenced without using long reads were mostly short (28). Furthermore, the collection of partial or repeat-domain-collapsed spidroin genes omitted at least one gene family. The new gene family of MaSp in *A. ventricosus* was found in a comparison of full-length sequences. Here, we used a hybrid method that combines the gDNA sequencing reads used for genome assembly and the a few hundred million reads obtained by cDNA and direct RNA sequencing to catalog the spidroin diversity in four Nephilinae spiders (S1 Table and Materials and Methods). Seven orthologous groups (MaSp: major ampullate spidroin, MiSp: minor ampullate spidroin, AcSp: aciniform spidroin, Flag: flagelliform spidroin, AgSp: aggregate spidroin, PySp: pyriform spidroin, and CySp: cylindrical/tubuliform spidroin) are known as the typical spidroins in Araneoid orb-weaving spiders. We obtained all seven groups contain 2-3 families or subfamilies from four Nephilinae draft genomes, five of which were full length (MaSp, MiSp, AcSp, CySp, and PySp) (Fig. 1, Fig. S5 for details).

**Figure 1.**
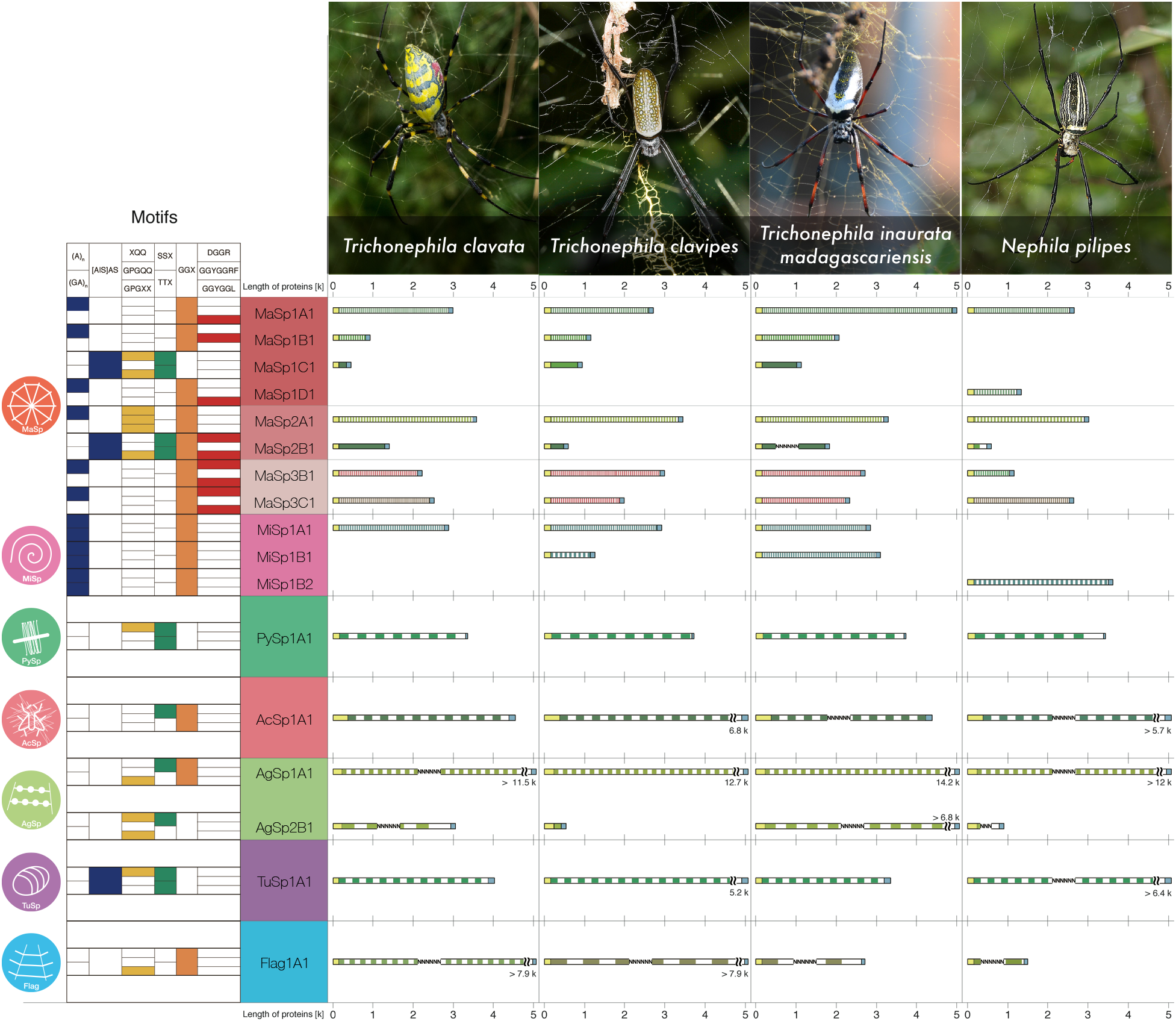
Catalog of spidroins in four Nephilinae spiders. This summary panel shows the spidroin sequence characteristics and structures obtained from four Nephilinae draft genomes. The icons in the first column represent spidroin groups. The second column represents the motif variety in the domains, as follows: β-sheet (An, (GA)n, and AS), blue; β-turn (XQQ, GPGQQ, and GPGXX), yellow; spacer (SSX and TTX), green; 310 helix (GGX), orange; and MaSp motif (DGGR, GGYGGRF, and GGYGGL), red. The spidroin structure columns show the N/C-terminal (yellow and blue box) and repeat domains, and each structure is drawn to scale. The number, width, and color of stripes reflect the number, size, and motif identify of the repeats. Assembly gaps are represented by “N” (see Fig. S5 for details).

The spidroin genes were well conserved among Nephilinae spiders and followed the established phylogenetic relationship (Fig. S6 and S7). Interestingly, the spidroin catalog revealed a group of major ampullate spidroins distinct from MaSp families 1 and 2 that form a unique clade within Nephilinae. The N-terminal phylogeny suggests that this group forms a clade with the MaSp family 3 sequence of Araneidae (Fig. S7), reflected by the characteristic DGGR motif, yet the N-terminus conservation is sufficiently low to define this as a unique group; thus, we named this gene MaSp3B based on a spidroin nomenclature (SI Text). Previously identified partial sequences (Sp-74867 and Sp-907) (27) belong to this group.

### Expression profile of spidroin and SpiCE

With this high-quality reference of full-length spidroins, the existence of a large proportion of MaSp3B in the forcibly reeled dragline silk of *T. clavata* was immediately observable as a distinct band in the high molecular weight (HMW) fraction on SDS-PAGE (Fig. 2*A*). The abundance ratios of the proteins were then quantified by intensity-based absolute quantification (iBAQ) (39) with shotgun proteomics of selectively reeled dragline silks (Fig. S9). These analyses showed consistent composition not only within the replicates but also among the four species analyzed (Fig. 2*B*). Among the spidroins, only MaSp family members were detected; MaSp1, 2, and 3 accounted for approximately 90 wt% of the total in any spider species (Fig. 2*B*). Transcriptome analysis of silk glands confirmed the specific expression of MaSp3B in the major ampullate silk gland, similar to that of MaSp1/2 (Fig. 2*C*, Fig. S10, and Table S2 to S4). Both of these omics analyses also clearly demonstrated the consistent presence of non-spidroins, which was also confirmed by SDS-PAGE of the LMW fraction (Fig. 2*A*). Across the four Nephilinae species, these genes are consistently expressed as mRNAs at an level equivalent to that of MaSp genes in the major ampullate silk gland and at lower levels in the minor ampullate silk gland (Fig 2C, Fig. S10). Three such proteins highly expressed in both silk (as protein) and the silk gland (as mRNA) were annotated as SpiCEs, following the identification of non-spidroin constituents in *A. ventricosus* dragline silk (28). These SpiCEs are conserved in Nephilinae (Fig. S11), but their sequences are considerably nonhomologous to those of *A. ventricosus*; therefore, we named them SpiCE-NMa1, 2, and 4 (SpiCE Nephilinae Major Ampullate). Among them, SpiCE-NMa2 and SpiCE-NMA4 were CRP type with high cysteine content. The most abundant non-spidroin protein among the dragline silks was SpiCE-NMa1, at approximately 1 wt% (Fig. 2*B*).

**Figure 2.**
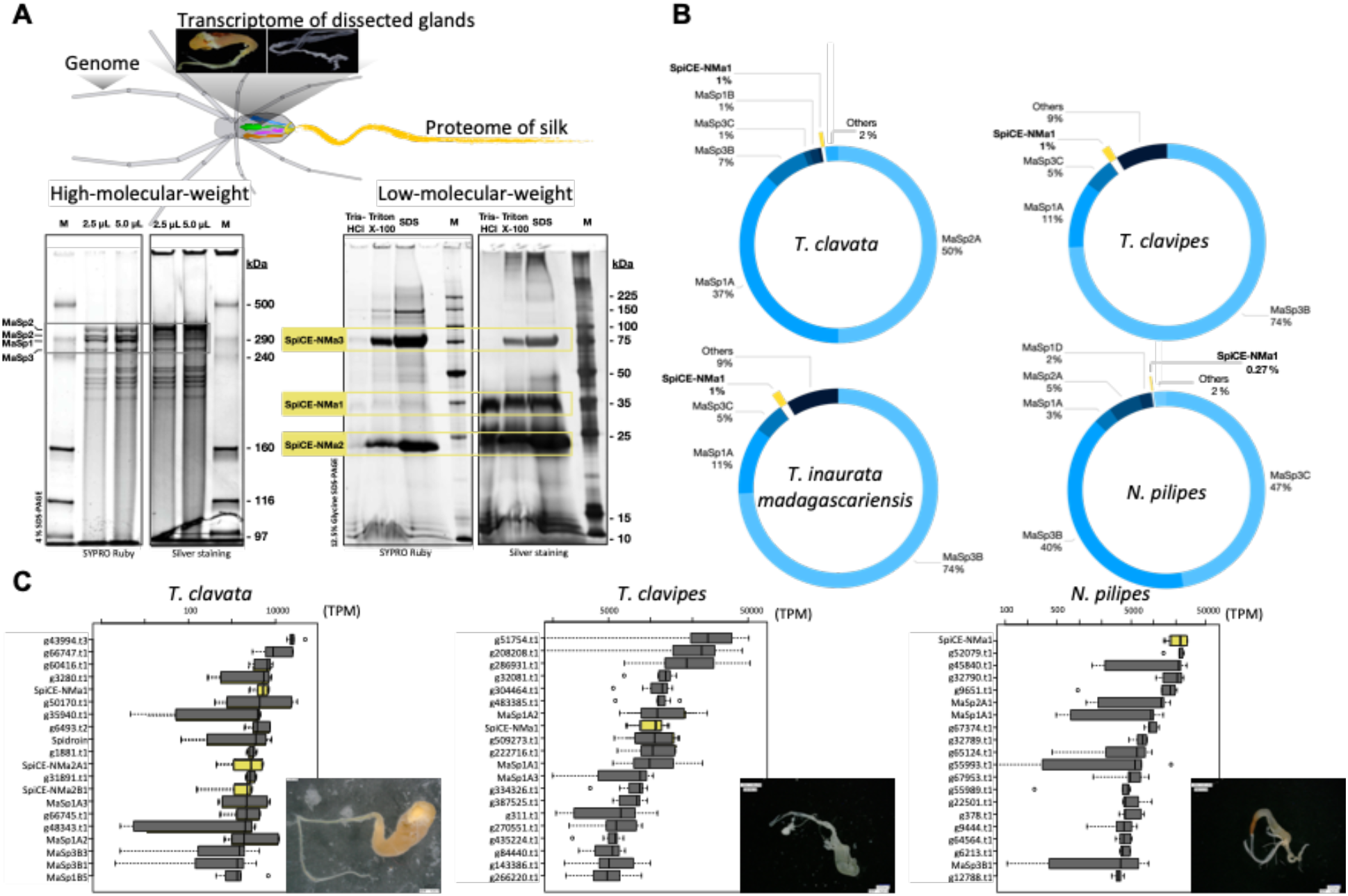
Expression profile of spidroin and SpiCE in silk and silk glands. (***A***) SDS-PAGE of HMW and LMW fractions from 3.97 mg of *T. clavata* dragline silk dissolved in 80 µL Tris-HCl (pH 8.6) containing 1% Triton-X and 2% SDS. SDS-polyacrylamide gels (4%) of the HMW fraction were stained with SYPRO Ruby or Silver after electrophoresis for 80 min at 20 mA. HiMark Unstained Standard (LC5688, Life Technologies) was used as a molecular marker. SDS-polyacrylamide gels (12.5%) of the LMW fraction were stained with SYPRO Ruby or Silver after electrophoresis for 80 min at 20 mA. Broad Range Protein Molecular Weight Marker (V8491, Promega Technologies) used as the molecular marker. (***B***) Average percentage of protein contents in dragline silk of *T. clavata*, *T. clavipes*, *T. inaurata madagascariensis*, and *N. pilipes*. Each SpiCE-NMa1 was indicated in yellow. (***C***) Expression profiles from the major ampullate silk glands in *T. clavata* (n = 6), *T. clavipes* (n = 7), and *N. pilipes* (n = 5). The x-axes represent the median expression level in TPM (transcripts per million). The y-axes show the highly expressed genes in each gland. The yellow boxes indicate SpiCE-NMa1. The photomicrographs show typical silk gland samples dissected from each spider (see S2 to S4 Table for details).

### SpiCE doubles the tensile strength of artificial silk-based material

To investigate the role of these proteins in the expression of the mechanical properties of Nephilinae dragline silks, we first produced composite films of recombinant MaSp family proteins. Recombinant MaSp was synthesized based on shortened amino acid sequences of approximately 50 kDa (Fig. S12 and S13). The transparency of silk-based films did not drastically decrease even when the composite films were prepared from two or three spidroins, suggesting that heterogeneous interactions among MaSps did not induce specific aggregation (Fig. 3*A*). Wide-angle X-ray scattering (WAXS) profiles demonstrated that MaSp1, MaSp2, and MaSp3 predominantly form β-sheet structures with different *d*-spacings based on the peak intensities at 6 and 18 nm^-1^ (Fig. 3*B*); however, the composite films of MaSps showed similar β-sheet structures. There were noticeable differences among the β-sheet structures with different protein compositions, that is, with MaSp1 used alone, with the mix of three families, and with the natural composition (Fig. 3*B*). We further tested the impact of SpiCE inclusion by producing composite films of recombinant MaSp and SpiCE-NMa1, which was approximately 5 times more abundant than other SpiCEs (Fig. 2*B*). Recombinant SpiCE-NMa1 was added in the range of 0 ∼ 5 wt% based on proteome quantification to produce composite films, and the film transparency increased to 76.1 with 5 wt% SpiCE-NMa1 with MaSp compared to 72.5 with MaSp alone, suggesting a close interaction of the proteins in the composite film (Fig. 4*A*, Fig. S14). Surprisingly, the addition of SpiCE-NMa1 dramatically changed the mechanical properties of the composite film. Tension tests showed a twofold increase in the tensile strength of the MaSp film containing SpiCE-NMa1 (Fig. 4, Fig. S15 and S16). The impact of SpiCE was also clearly observed in the recombinant composite silk mixed with recombinant MaSp and SpiCE-NMa1. Composite silk with SpiCE-NMa1 showed an increase in elongation at break, and notably, the stress-strain curve showed a clear drastic yield point only in the composite silk mixed with SpiCE-NMa1 (Fig. 4*C*). Natural spider silk is characterized by a nonlinear response to external pressure, and softening at the yield point allows for the mechanical robustness of the spider web (40–44). Therefore, including a small percentage (∼2%) of SpiCE-NMa1 in artificial silks could potentially be used to enhance their mechanical properties.

**Figure 3.**
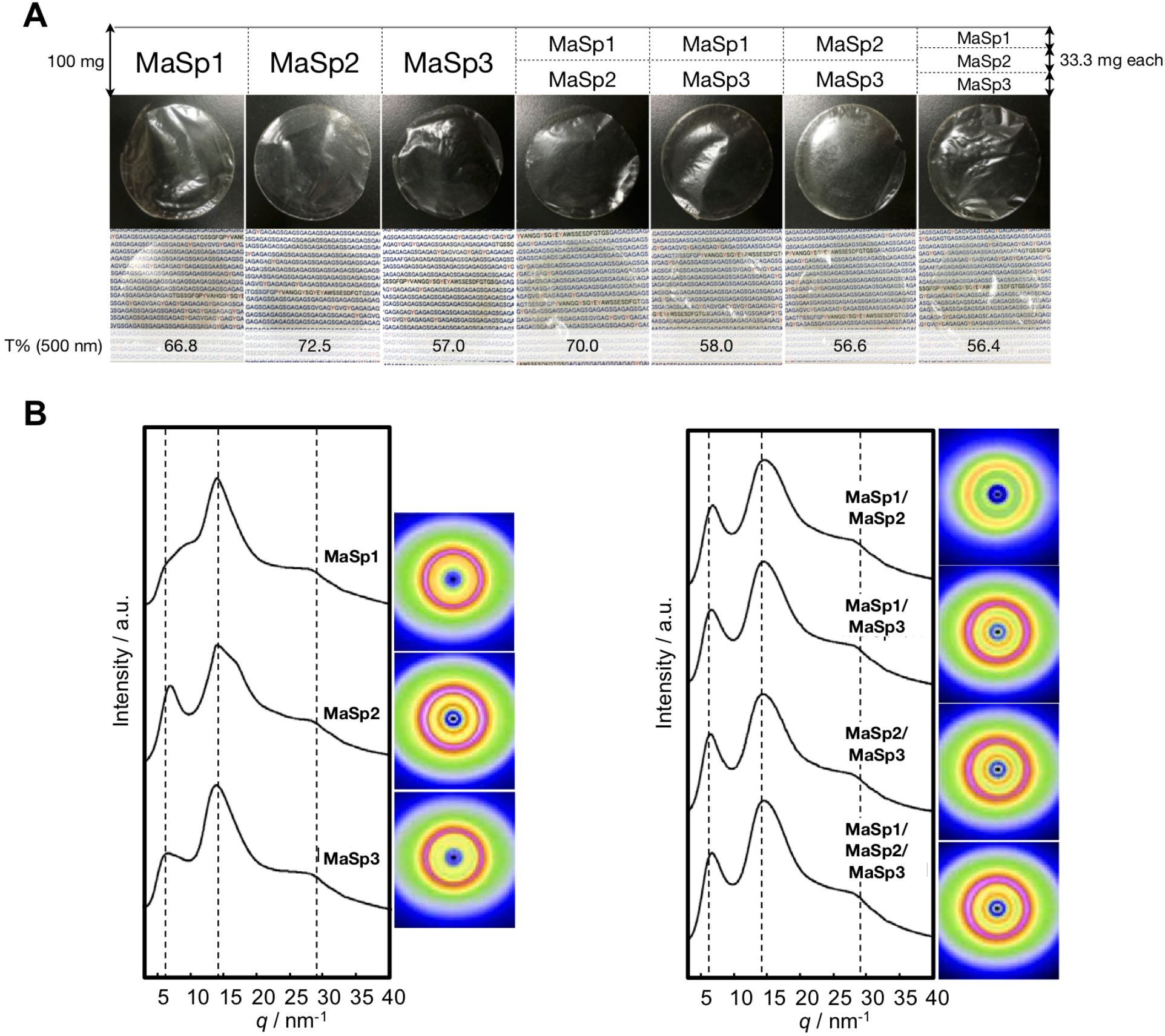
Physical properties of the artificial MaSp composite film. (***A***) Seven films were produced by the combination of recombinant MaSp family proteins (MaSp1, MaSp2, and MaSp3). The sequences of recombinant proteins are indicated in Fig. S11. T% indicates transparency at a wavelength of 500 nm. (***B***) The WAXS profiles of the films with different MaSp compositions showing the one-dimensional radial integration profiles and two-dimensional patterns.

**Figure 4.**
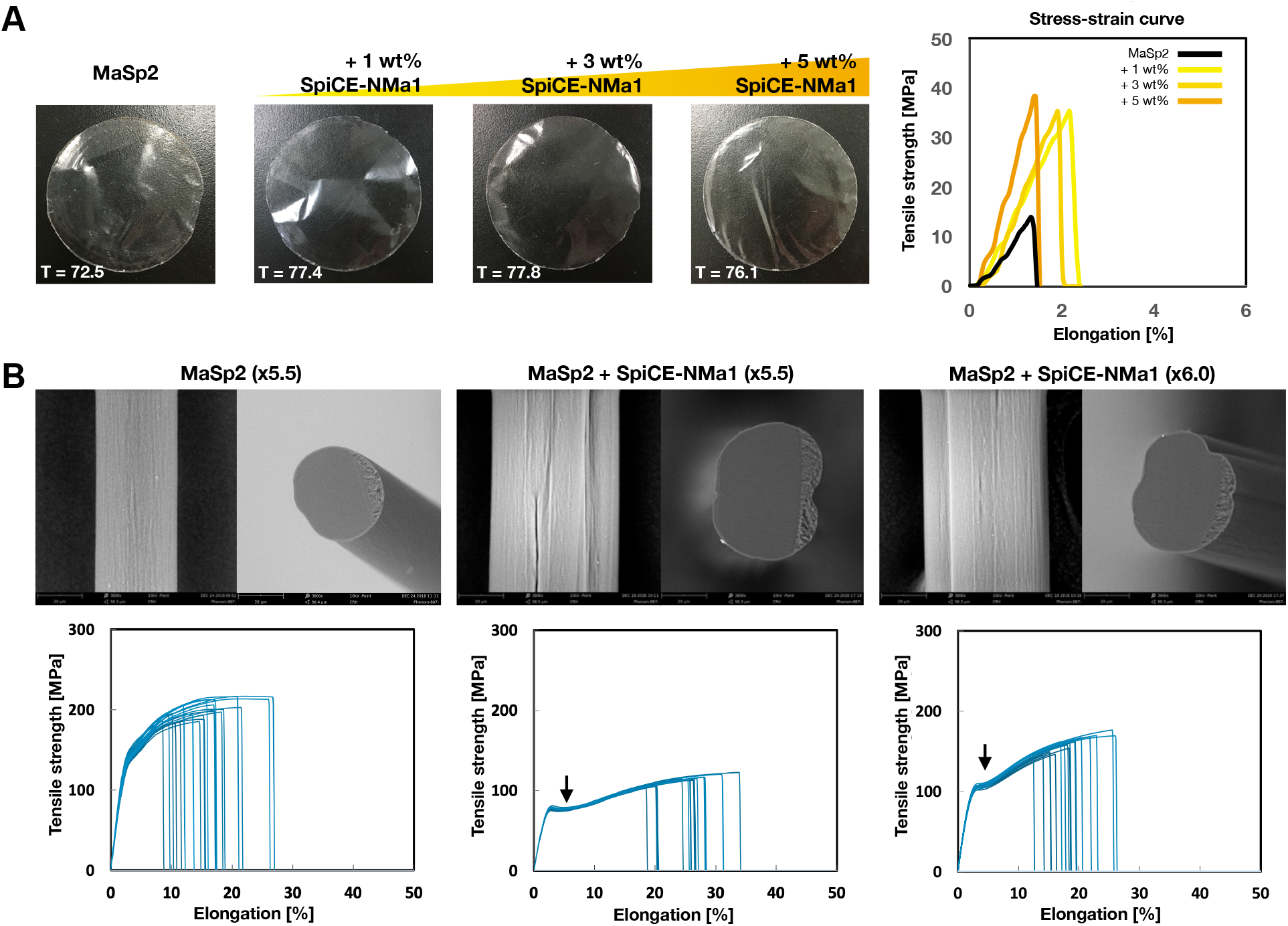
SpiCEd film and silk. (***A***) Composite films mixing MaSp2 and SpiCE-NMa1. The stress-strain curve represents the typical tensile engineering stress-strain in MaSp2 and SpiCE composite films under different SpiCE concentrations (0, 1, 3, 5 wt%). The transparency (%) of these films is shown as T in each picture (see Fig. S13-S15 for details). (***B***) Recombinant spider silks mixing MaSp2 and SpiCE-NMa1 with different draw ratios (5.5 or 6.0). Scanning electron microscopy (SEM) images show the cylindrical and smooth surface and cross-section of each recombinant spider silk. (***C***) Stress-strain curve of each recombinant spider silk. Black arrows indicate the yield points.

## Discussion

MaSp3 is widely conserved in the large-web-forming group of the family Araneidae and has been found by proteomics to be a major component of dragline silk in *A. ventricosus*. The conservation of MaSp3 has been thought to be limited to closely related species of the genus *Araneus* (28), and it was believed that Nephilinae spiders do not have MaSp3 even though they also build large webs. However, our high-quality genome analyses demonstrate that MaSp3B, a distinct subtype of MaSp3 in the genus *Araneus*, is conserved in the genera *Trichonephila* and *Nephila.* Further, our analyses suggest that MaSp3B is one of the major components of dragline silk and may associate with other MaSps at equivalent stoichiometry. This finding demonstrates the limitation of the conventional MaSp1/2-based strategy for artificial silk synthesis and the importance of including other components, such as MaSp3, for the purpose of producing araneoid spider silk, which has particularly strong mechanical properties. Although the specific role of MaSp3 is still unclear, the key residues contributing to the N-terminal domain association, such as D40, K65, E79, and E119, are conserved in MaSp1-3 (Fig. S17). Therefore, it is possible that the formation of a heteromer with MaSp1/2 may stabilize the silk structure and improve its toughness.

The contribution of the low molecular weight component of the spider dragline silk, SpiCE, to its mechanical properties was also confirmed in vitro, in that inclusion of SpiCE doubled the tensile strength of the artificial film, but the exact mechanism responsible for this phenomenon is still an area of future work. Accessory components of structural proteins are often used to stabilize the structure; for example, elastin, another known structural protein, binds with fibulin proteins to produce elastic fibers (45). Likewise, while a self-assembly system involving pH-induced fibrillization of the N-terminal domain and liquid-liquid phase separation (LLPS) of the C-terminal domain (14) is known to participate in spider silk fibril formation, SpiCE could serve to support the interaction between repeat regions rather than the terminal domains. Further observation of the interaction between SpiCE and spidroin with detailed biophysical analysis, such as nuclear magnetic resonance (NMR) spectroscopy or atomic force microscopy (AFM) scanning, is desirable to elucidate the functions of SpiCE. The evolutionary origin of SpiCE also requires further analyses, as these proteins seem to be specific to a narrow range of spider clades. Unlike MaSp3, the conservation of SpiCE has been confirmed only within very closely related groups (genus or subfamily at most), and CRP seems to be used instead of SpiCE in black widow spiders (26). The MaSps retain homology across different spider clades, but there is no sequence homology among SpiCEs, suggesting that MaSps and SpiCEs did not coevolve. The orb web size evolved in response to body size, environment, and prey, and the dragline silk made by MaSp used for the orb web frame needed to be strong enough to match its size (46). To produce silks with stronger properties, mutations in the nonconserved repeat domain of the spidroin gene and diversification of multiple MaSp families by gene duplication must have occurred. In fact, multiple lineage-specific duplications of MaSp paralogs are observed even among orb-weaving spiders (27, 28, 47-49). The evolution of SpiCE seems to follow this lineage-specific evolutionary pattern.

Considering the contributions of MaSp3 and SpiCE to the mechanical performance of spider silk, it is essential to consider silk as not just a product of MaSp1 and MaSp2 but rather a multicomponent material that also includes MaSp3 and SpiCE. The contribution of multiple paralogs of MaSp1 and MaSp2, which is not covered in this work, may also contribute to the specific ecological adaptation of *Nephilinae* spiders. The availability of four closely related Nephilinae genomes and their gland transcriptomes as well as the dragline silk proteomes presented in this work will contribute to the research in this direction, as well as other phylogenomic work of arachnids.

## Materials and Methods

### Spider sample and rearing environment

*Trichonephila clavata* and *Nephila pilipes* samples were collected in Japan, and *Trichonephila inaurata madagascariensis* was collected in Madagascar. *Trichonephila clavipes* samples were purchased from Spider Pharm Inc. All the spiders used in this study were adult females. The identification of spider specimens was performed based on morphological characteristics and sequence identification of cytochrome c oxidase subunit 1 (*COX1*) in the Barcode of Life Data System (BOLD: http://barcodinglife.org). The spiders used for dissection or silk reeling were kept in a PET cup (129 x 97 mm, PAPM340: RISUPACK CO., LTD) in a climate-controlled laboratory with 25.1 °C, 57.8% humidity and 12-12 h light-dark cycles. The spiders were fed every two days with crickets purchased from Mito-korogi Farm, and water was given daily by softly spraying inside the cup. The legs and cephalothoraxes were used for gDNA extraction, and abdomens and silk glands were used for RNA extraction. All of the tissues were dissected from adult female spiders. Dissected samples were immersed in liquid nitrogen (LN2) and stored at −80 °C until the next process.

### High molecular weight genomic DNA extraction

Spider genomic DNA was extracted from dissected leg and cephalothorax tissue from adult female individuals using Genomic-tip 20/G (QIAGEN) following the manufacturer’s protocol. To obtain high molecular weight (HMW) genomic DNA, all steps were conducted as gently as possible. The legs and cephalothorax were separated from flash-frozen spider specimens and then homogenized with a BioMasher II (Funakoshi) and mixed with 2 mL of Buffer G2 (QIAGEN) containing 200 µg/mL RNase A. After the addition of 50 µL Proteinase K (20 mg/mL), the lysate was incubated at 50 °C for 12 h on a shaker (300 rpm) and centrifuged at 5,000 x g for 5 min at 4 °C. The aqueous phase was loaded onto a pre-equilibrated QIAGEN Genomic-tip 20/G (QIAGEN) by gravity flow and washed three times. The DNA was eluted in high-salt buffer (Buffer QF) (QIAGEN). Using isopropanol precipitation, we desalted and concentrated the eluted DNA and resuspended it in 10 mM Tris-HCl (pH 8.5). The extracted genomic DNA was checked for quality using a TapeStation 2200 instrument with genomic DNA Screen Tape (Agilent Technologies) and quantified using a Qubit Broad Range dsDNA assay (Life Technologies). Size selection (> 10 kb) was performed with a BluePippin with High Pass Plus Gel Cassette (Sage Science).

### RNA extraction, mRNA selection, and cDNA preparation

Spiders and silk gland samples were stored in LN2 at −80 °C until RNA extraction. RNA isolation from the dissected abdomen or silk glands in the abdomen was conducted based on a spider transcriptome protocol (50). Flash-frozen dissected abdomen tissue was immersed in 1 mL TRIzol Reagent (Invitrogen) along with a metal cone and homogenized with the Multi-Beads Shocker (Yasui Kikai). Phase separation was performed by the addition of chloroform, and the upper aqueous phase containing RNA was purified automatically with an RNeasy Plus Mini Kit (QIAGEN) in a QIAcube instrument (QIAGEN). The RNA was quantified and checked for quality using a Qubit Broad Range RNA assay (Life Technologies) and NanoDrop 2000 (Thermo Fisher Scientific). The integrity was estimated by electrophoresis using a TapeStation 2200 instrument with RNA ScreenTape (Agilent Technologies). mRNA selection was performed using oligo d(T). For direct RNA sequencing, mRNA was prepared using the NucleoTrap mRNA Mini Kit (Clontech) from 500 ng of total RNA. cDNA was synthesized from mRNA isolated from 100 µg of total RNA by NEBNext Oligo d(T)_25_ beads (skipping the Tris buffer wash step). The first- and second-strand cDNA were synthesized using ProtoScript II Reverse Transcriptase and NEBNext Second Strand Synthesis Enzyme Mix.

### Library preparation and sequencing

For cDNA sequencing was carried out on the abdomens of *T. clavata* (n = 14), *T. clavipes* (n = 2), *T. inaurata madagascariensis* (n = 1) and *N. pilipes* (n = 14); in the major ampullate silk glands of *T. clavata* (n = 6), *T. clavipes* (n = 7), and *N. pilipes* (n = 5); and in the minor ampullate silk glands of *T. clavata* (n = 6), *T. clavipes* (n = 8), and *N. pilipes* (n = 6). All cDNA libraries were constructed according to the standard protocol of the NEBNext Ultra RNA Library Prep Kit for Illumina (New England BioLabs). The synthesized double-stranded cDNA was end-repaired using NEBNext End Prep Enzyme Mix before ligation with NEBNext Adaptor for Illumina. After USER enzyme treatment, cDNA was amplified by PCR with the following conditions: 20 µL cDNA, 2.5 µL Index Primer, 2.5 µL Universal PCR Primer, 25 µL NEBNext Q5 Hot Start HiFi PCR Master Mix 2X; 98 °C for 30 s and 12 cycles each of 98 °C for 10 s, 65 °C for 75 s and 65 °C for 5 min. cDNA sequencing was conducted with a NextSeq 500 instrument (Illumina) using 150-bp paired-end reads with a NextSeq 500 High Output Kit (300 cycles).

The synthetic long-read sequencing using 10X Genomics was carried out in all four Nephilinae spiders. Purified genomic DNA fragments longer than 60 kb (10 ng) were used to prepare the 10X GemCode library with the Chromium instrument and Genome Reagent Kit v2 (10X Genomics) following the manufacturer’s protocol. The 10X GemCode library sequencing was conducted with a NextSeq 500 instrument (Illumina) using 150-bp paired-end reads with a NextSeq 500 High Output Kit (300 cycles).

For PacBio sequencing, the library preparation was performed with gDNA fragments of more than 10 kb in size (selected by BluePippin) and sequenced in PacBio RSII using P6-C4 chemistry. PacBio library construction and sequencing were conducted at Genomic Information Research Center, Osaka University.

The gDNAs of *T. clavata*, *T. clavipes*, *T. inaurata madagascariensis*, and *N. pilipes* were also sequenced by Nanopore technology. Library preparation was completed following the 1D library protocol (SQK-LSK109, Oxford Nanopore Technologies). The quality of the libraries was estimated by TapeStation 2200 with D1000 Screen Tape (Agilent Technologies). Direct RNA sequencing was performed for *T. clavata*, *T. clavipes*, and *N. pilipes*. Libraries constructed from 500 ng of total RNA were prepared following the direct RNA sequencing protocol (SQK-RNA001, Oxford Nanopore Technologies). Sequencing was performed using a GridION instrument with Spot On Flow Cell Rev D (FLO-MIN106D, Oxford Nanopore Technologies). The base calls were performed by Guppy basecalling software (version 3.2.10+aabd4ec).

### Genome assembly

We adopted a hybrid assembly strategy for spider draft genomes that combined the various reads produced by each sequencing technology. Natural long reads were produced by direct sequencing of HMW genomic DNA with Nanopore or PacBio technology. The synthetic long reads were generated using a combination of Illumina and 10X Genomics technologies. The details of the assembly methods used for each of the four spider genomes are described below.

#### T. clavata

Long reads produced by PacBio RSII were assembled with Canu 1.4 (51) with default parameters. First, polishing was performed with Quiver in SMRT analysis 2.3.0 and further polished with pilon 1.22 (52). Synthetic long reads of the 10X GemCode library sequenced by a NextSeq 500 instrument were assembled by Supernova 2.1 with pseudohap2 output mode (53). The above two assemblies were merged using SSPACE-LongRead v.1.1 with the parameters (-i 95 -o 2000 -l 1 -k 1) (54).

#### T. clavipes

Long reads from Nanopore sequencing were first quality filtered using NanoFilt with options -q 7 --headcrop 50 (55), and then assembled using Flye 2.7 with three polishing iterations (56). The assembly was further polished by pilon 1.23 (52) using 10X GemCode library sequencing reads for four rounds until the BUSCO score was saturated.

#### T. inaurata madagascariensis

Synthetic long reads of the 10X GemCode library sequenced by a NextSeq 500 instrument were assembled by Supernova 2.1.1 with pseudohap2 output mode (53). Assembled contigs were gap-filled using PBJelly (support option -m 45, blasr option -minMatch 8 -minPctIdentity 70 - bestn 1 -nCandidates 20 -maxScore -500 -nproc 32 -noSplitSubreads) (57) and polished by pilon v.1.23 (52) for three rounds. Long reads from Nanopore sequencing were first quality filtered using NanoFilt with options -q 7 --headcrop 50 (55), assembled using wtdbg v2.0 (58), and then further polished by pilon v.1.23 for three rounds. Two assemblies were merged using quickmerge with options -hco 5.0 -c 1.5 -l 100000 - ml 5000 (59).

#### N. pilipes

Synthetic long reads from the 10X GemCode library sequenced by a NextSeq 500 instrument were assembled by Supernova 2.0.0 with pseudohap2 output mode (53). The resulting assemblies were further merged and gap-filled using PBJelly (blasr option -minMatch 8 -minPctIdentity 70 -bestn 1 - nCandidates 20 -maxScore -500 -nproc 32 -noSplitSubreads) (57) and polished by pilon 1.22 (52).

### Contaminant elimination and genome size estimation

To detect possible contaminants in the genome, we submitted each genome assembly to BlobTools analysis (60). The genome sequence was submitted to a Diamond (61) BLASTX search (--sensitive --max-target-seqs 1 --evalue 1e-25) against the UniProt Reference proteome database (downloaded on 2018 Nov.). Each contig was classified into a lineage with BlobTools taxify. We then mapped DNA-Seq reads to the genome by BWA MEM and conducted SAM to BAM conversion with SAMtools. These taxonomic annotation and coverage data were visualized by BlobTools following the protocol. Contigs that were classified as from bacteria, plants, or fungi were removed from the assembly, and the filtered genome assembly was assayed by Benchmarking Universal Single-Copy Orthologs (BUSCO) v4.0.5 (33) (eukaryote lineage) to validate genome completeness. Genome heterozygosity, repeat content, and size were estimated based on the k-mer distribution with Jellyfish and GenomeScope (62).

### Gene prediction and annotation

Gene prediction was performed using a gene model created by cDNA-seq mapping data with HISAT2 and BRAKER (63, 64). cDNA-seq reads were mapped to the genome with HISAT2 (v 2.1.0), and the resulting SAM file was converted, sorted, and indexed by SAMtools (v1.4). Repeat sequences were detected by RepeatModeler (1.0.11) and soft-masked by RepeatMasker (v4.0.7). The soft-masked genome was submitted to gene prediction with BRAKER (v2.1.4, --softmasking --gff3). The amino acid sequences were submitted to BLASTP or Diamond BLASTP searches against public databases (UniProt TrEMBL, UniProt Swiss-Prot). Redundant genes were eliminated by CD-HIT-EST (35) clustering with a nucleotide identity of 97%. To obtain a functional gene set, we removed the genes with an expression level of less than 0.1 and unannotated genes. BUSCO (v4.0.5) was used to determine the quality of our functional gene set using the eukaryote lineage. The tRNAs were annotated using tRNAscan-SE v2.0 (34) with default parameters. Ribosomal RNAs (rRNAs) were predicted using Barrnap (https://github.com/tseemann/barrnap).

### Spidroin gene catalog

We used a hybrid method that combines short- and long-read sequencing to catalog the spidroin diversity in four Nephilinae spiders. The short reads were obtained from Illumina sequencing of cDNA or gDNA libraries, and the long reads were obtained by Nanopore or PacBio sequencing of the gDNA library. The typically used de Bruijn graph algorithm is not suitable for the assembly of long repeat domains, so the SMoC (Spidroin Motif Collection) algorithm (28), developed based on the OLC (Overlap-Layout-Consensus) algorithm, was used. The SMoC algorithm collects as many patterns of repeat motifs as possible, scaffolds these repeat motifs based on long reads, and then provides the full-length spidroin gene sequence. SMoC first finds the N/C-terminal region (nonrepetitive region) contigs from the assembled contigs with a homology search. These terminal fragments were used as seeds for screening of the short reads obtained by an Illumina sequencer harboring an exact match of extremely large k-mers (approximately 100) up to the 5′-end. The obtained short reads were aligned on the seed sequence to construct a PWM (position weight matrix) on the 3′-side of the matching k-mer, and the seed sequence was extended based on the PWM until there was a split in the graph using stringent thresholds. Therefore, neighboring repeats are not resolvable. After the comprehensive collection of repeat motif patterns, scaffolding was performed using long reads. The long reads were obtained by direct sequencing, so the actual gene length was guaranteed. Finally, mapping the repeat motifs to the corresponding long read and curating the sequence yields the complete spidroin genes.

### Phylogenetic tree of spidroin

The phylogenetic tree in Fig. S6 was constructed based on a core ortholog gene set obtained from BUSCO arthropoda_obd9 (33) using four Nephilinae draft genomes (this study) and the *A. ventricosus* draft genome (Ave_3.0) (28). The phylogenetic tree of spidroin (Fig. S7) was constructed by FastTree (v2.1.10) (65) as an approximately maximum-likelihood phylogenetic tree from aligned and trimmed N-terminal sequences (150 residues from the start codon) with MAFFT (v7.273) (66) and trimAl (1.2rev59) (67).

Accession numbers for known spidroin sequences are as follows: major ampullate spidroin 2-like (AAZ15322.1), and flagelliform silk protein (AAF36091.1) in *T. inaurata madagascariensis*; major ampullate spidroin 1A precursor, partial (ACF19411.1), major ampullate spidroin 1B precursor, partial (ACF19412.1), major ampullate spidroin 2 precursor, partial (ACF19413.1), major ampullate spidroin protein MaSp-a (PRD23950.1), major ampullate spidroin protein MaSp-b (PRD18716.1), major ampullate spidroin protein MaSp-c (PRD18936.1), major ampullate spidroin protein MaSp-d (PRD23750.1), major ampullate spidroin protein MaSp-f isoform 1 (PRD27696.1), major ampullate spidroin protein MaSp-g (PRD24320.1), major ampullate spidroin protein MaSp-h (PRD20448.1), minor ampullate spidroin protein MiSp-a (PRD23654.1), minor ampullate spidroin protein MiSp-b (PRD30268.1), minor ampullate spidroin protein MiSp-c (PRD24510.1), minor ampullate spidroin protein MiSp-d (PRD18914.1), flagelliform silk protein (AAC38846.1), aciniform spidroin protein AcSp (PRD26201.1), aggregate spidroin protein AgSp-a (PRD23399.1), aggregate spidroin protein AgSp-b (PRD22063.1), aggregate spidroin protein AgSp-c (PRD26655.1), aggregate spidroin protein AgSp-d (PRD23989.1), tubuliform spidroin protein TuSp (PRD35275.1), flagelliform spidroin protein FLAG-a (PRD27227.1), flagelliform spidroin protein FLAG-b (PRD24772.1), piriform spidroin protein PiSp (PRD25616.1), and spidroin protein Sp-907 (PRD35000.1), and spidroin protein Sp-74867 (PRD19552.1) in *T. clavipes*; cylindrical silk protein 1 (BAE54451.1) in *T. clavate*; major ampullate spidroin 3 variant 1, partial (AWK58729.1) in *Argiope argentata*; Major ampullate spidroin 3 (GBN25680.1) in *A. ventricosus*; aggregate spidroin 1 (QDI78451.1), aggregate spidroin 2 (QDI78449.1) in *Argiope trifasciata*; eggcase silk protein (ACI23395.1) in *Nephila antipodiana*; egg case silk protein 1 (BAE86855.1) in *Argiope bruennichi*.

### Gene expression analysis

Gene expression profiling was conducted with multiple biological replicates from mRNA extracted from the abdomen or silk glands. The gene expression levels were quantified and normalized as transcripts per million (TPM) by mapping processed reads to our assembled draft genome references with Kallisto version 0.42.1 (68) (S2 to S4 Table).

### Silk collection

The natural spider silks were sampled directly from adult female *T. clavata* restrained using two pieces of sponge and locked with rubber bands. Silk reeling from the spider was performed at a constant speed (1.28 m/m for 1 h) with a reeling machine developed by Spiber Inc. with six biological replicates and three technical replicates each. Since the spinnerets are very close together and many silks may be spun at the same time, simple silk reeling may involve minor ampullate silk as well. Therefore, to identify the proteins that were significantly present in minor ampullate silk and remove them as background, we separated the dragline and minor ampullate silks clearly from their spinnerets using a stereomicroscope (Leica S4E, Leica microsystems GmbH, Wetzlar, Germany) (Fig. S9).

### SDS-PAGE analysis

SDS-PAGE analysis of dragline silk was conducted at IDEA Consultants, Inc. The dragline silk reeled from *T. clavate* was dissolved in 2 mL of ionic liquid (1-butyl-3-methylimidazolium acetate) per mg. After vortexing for 2 min, the sample was incubated at 100 °C for 15 min. The ionic liquid solution was dialyzed into lysis buffer (6 M urea, 2 M thiourea, 2% CHAPS, 1% DTT) using a 3k MWCO cellulose membrane (Amicon Ultra, Millipore).

The high molecular weight fraction of 0.33 mg dragline silk was dissolved in the ionic liquid and filtered through a 50k MWCO membrane (Amicon Ultra, Millipore), collected in 100 µL of lysis buffer, and incubated at 95 °C for 5 min. Then, 2.5 µL and 5 µL samples were applied to a 4% SDS-polyacrylamide gel. Electrophoresis was performed at 20 mA for 80 min in running buffer [25 mM Tris, 192 mM glycine, 0.1% SDS], and a HiMark Unstained Standard (LC5688, Life Technologies) was used as a molecular marker. After electrophoresis, the SDS gel was fixed in fixing solution [30% MeOH, 5% acetate acid], stained with SYPRO Ruby overnight, and washed with a wash solution [10% MeOH, 7% acetic acid]. Silver staining was conducted after fixation with 50% MeOH and sensitization with 0.005% sodium thiosulfate (Fig. 2*A*).

A solution of 3.97 mg dragline silk dissolved in 80 µL of Tris-HCl (pH 8.6), 1% Triton-X, and 2% SDS was vortexed, incubated for 1 h on ice, and freeze-dried. After resuspension, the total volume was applied to a 12.5% SDS-polyacrylamide gel. Electrophoresis was performed at 20 mA for 80 min, with Broad Range Protein Molecular Weight Markers (V8491, Promega) used as molecular markers. After electrophoresis, the SDS gel was fixed in fixing solution [30% MeOH, 5% acetic acid], stained with SYPRO Ruby overnight, and washed with a wash solution [10% MeOH, 7% acetic acid]. Silver staining was conducted after fixation with 50% MeOH and sensitization with 0.005% sodium thiosulfate (Fig. 2*A*).

### Liquid chromatography mass spectrometry (LC-MS) analysis

Approximately 1.0 mg of dragline silk was used for the LC-MS analysis. After washing the *T. clavipes* or *N. pilipes* dragline silk with 100 µL of washing buffer [50 mM NH_4_HCO_3_, 0.1% SDS], the silk was immersed in 50 µL of solution A [50 mM NH_4_HCO_3_, 500 mM DTT] and incubated at 60 °C for 1 h. The silk was further incubated with 50 µL of solution B [50 mM NH_4_HCO_3_, 500 mM IAA] at room temperature for 30 min in the dark and washed with 100 µL of 50 mM NH_4_HCO_3_. Silk protein digestion was carried out in 50 µL of digestion buffer [10 ng/µL trypsin, 50 mM NH_4_HCO_3_] at 37 °C overnight. The digested sample was incubated with 250 µL of 0.5% formic acid on a rotator and purified using a MonoSpin C18 column (GL Science).

The dragline silk of *T. clavata* or *T. inaurata madagascariensis* was immersed in 200 µL of lysis buffer [6 M guanidine-HCl, pH 8.5] per mg dragline silk and frozen in liquid nitrogen. After 10 cycles of sonication with 60 sec on/off at high level using Bioruptor II (BM Equipment), the protein was quantified with Qubit Protein assay (Life Technologies) and Pierce BCA Protein Assay Kit (Thermo Scientific). The lysate [50 µg protein/50 µL] was incubated with 0.5 µL of 1 M DTT solution at 37 °C for 30 min followed by 2.5 µL of 1 M IAA solution at 37 °C for 30 min in the dark. After five-fold dilution with 50 mM NH_4_HCO_3_, protein digestion was performed by 1 µg of Lys-C (Wako) at 37 °C for 3 h following 1 µg of trypsin (Promega, Madison, WI, USA) at 37 °C for 16 h. The sample was acidified by TFA and desalted by C18-StageTips (69). LC-MS analysis of *T. clavipes* (n = 6) and *N. pilipes* (n = 9) samples was performed with an UltiMate 3000 NanoLC Pump (Dionex Co., Sunnyvale, CA, USA) and an LTQ Orbitrap XL ETD (Thermo Electron, San Jose, CA, USA). A 5 µg aliquot of digested sample was injected into a spray needle column [ReproSil-Pur C18-AQ, 3 µm, Dr. Maisch, Ammerbuch-Entringen, Germany, 100 µm i.d.×130 mm] and separated by linear gradient elution with three mobile phases, A [0.5% acetic acid in water], B [0.5% acetic acid in acetonitrile] and C [0.5% acetic acid in DMSO], at a flow rate of 500 nL/min. The composition of mobile phase C was kept at 4%. The composition of mobile phase B was changed from 0-4% in 5 min, 4-24% in 60 min and 24-76% in 5 min, followed by keeping at 76% in 10 min. The separated peptides were ionized at 2600 V and detected as peptide ions (scan range: m/z 300-1500, mass resolution: 60000 at m/z 400). The top 10 peaks of multiple charged peptide ions were subjected to collision-induced dissociation (isolation width: 2, normalized collision energy: 35 V, activation Q: 0.25, activation time: 30 s).

The samples of *T. clavata* (n = 7) and *T. inaurata madagascariensis* (n = 1) were analyzed with a nanoElute and a timsTOF Pro (Bruker Daltonics, Bremen, Germany). A 200 ng aliquot of digested sample was injected into a spray needle column [ACQUITY UPLC BEH C18, 1.7 µm, Waters, Milford, MA, 75 µm i.d.×250 mm] and separated by linear gradient elution with two mobile phases, D [0.1% formic acid in water] and E [0.1% formic acid in acetonitrile], at a flow rate of 280 nL/min. The composition of mobile phase E was increased from 2% to 35% in 100 min, changed from 35% to 80% in 10 min and kept at 80% for 10 min. The separated peptides were ionized at 1600 V and analyzed by parallel accumulation serial fragmentation (PASEF) scan (70). Briefly, the PASEF scan was performed at ion mobility coefficients (1/K0) ranging from 0.6 Vs/cm^2^ to 1.6 Vs/cm^2^ within a ramp time of 100 msec, keeping the duty cycle at 100%. An MS scan was performed in the mass range from m/z 100 to m/z 1700, followed by 8 PASEF-MS/MS scans per cycle. Precursor ions were selected from the 12 most intense ions in a TIMS-MS survey scan (precursor ion charge: 0-5, intensity threshold: 1250, target intensity: 10000). In addition, a polygon filter was applied to the m/z and ion mobility plane to select the most likely representative peptide precursors without singly charged ions. CID was performed with the default settings (isolation width: 2 Th at m/z 700 and 3 Th at m/z 800, collision energy: 20 eV at 1/k0 0.6 Vs/cm^2^ and 59 eV at 1/k0 1.6 Vs/cm^2^). LC-MS data of *T. clavipes* (n = 6) and *N. pilipes* (n = 9) samples were analyzed using PEAKS studio X+ (Bioinformatics Solutions Inc., Canada) with the following conditions (71). Briefly, de novo sequencing and database searches were performed with an error tolerance of 6 ppm for precursor ions and 0.5 Da for fragment ions. The enzyme was set to trypsin, and up to 2 missed cleavages were allowed. Carbamidomethylation at cysteine residues was set as a fixed modification. N-acetylation at the protein N-terminus and oxidation at the methionine residue were set as variable modifications, allowing for up to 3 positions per peptide. A protein sequence database generated from our draft genome was used for identification. The MaxQuant (version 1.6.10.43) (72) contaminant database (245 entries, major experimental contaminants) was used to detect contaminants. The criterion of identification was set to less than 1% FDR at the peptide-spectrum match (PSM) level. The feature area of each identified peptide ion was calculated automatically with the PEAKS software algorithm. The intensity-based absolute quantification (iBAQ) (39) value of each identified protein was calculated from the feature area values. LC-MS data of *T. clavata* (n = 7) and *T. inaurata madagascariensis* (n = 1) were also analyzed using the same PEAKS software. De novo sequencing and database search were performed with an error tolerance of 20 ppm for precursor ions and 0.05 Da for fragment ions. The protein sequence database obtained from our draft genome was used for protein identification. Other conditions were the same as described above.

To eliminate the possibility that other silks might have contaminated reeled dragline silk samples, we removed background data. We separated the dragline and minor ampullate silks clearly from the spinneret by microscopy and used them for LC-MS analysis to identify the background proteins. Using the iBAQ score, we identified the proteins specifically expressed in the minor ampullate silk based on significance (FDR < 0.01) and removed them from the result of the dragline silk proteome analysis as contaminants. The percentage of each protein in silk was calculated based on the iBAQ multiplied by the amino acid length.

### Recombinant protein expression and purification

Sequence-optimized recombinant *T. clavata* MaSp family and SpiCE-NMa1 proteins were used for the composite film and silk. Targeting a size of approximately 50 kDa, we reduced the number of repeat units to 14 for MaSp1, 9 for MaSp2, and 8 for MaSp3 while keeping the N- and C-terminal domains (Fig. S12). The recombinant spidroin genes are composed of a 6x His tag (MHHHHHH), a linker (SSGSS), an HRV 3C protease recognition site (LEVLFQGP), an N-terminal domain, repeat units, and a C-terminal domain. The *SpiCE-NMa1* gene is also composed of a 6x His tag, a linker (SSGSS), an HRV 3C protease recognition site and a *T. clavata* comp1999 protein sequence. Each signal peptide was removed based on SignalP-5.0 (73). Nucleotide sequences were adjusted to the codon usage of *E. coli*. Fragmented sequences were chemically synthesized by Fasmac Co., Ltd. (Atsugi, Japan). The synthesized sequences were assembled with overlap extension PCR (74). Assembled sequences were cloned into pET-22b(+) and transformed into *E. coli* BLR (DE3). A 100 mL bacterial culture (5.0 g/L glucose, 4.0 g/L KH_2_PO_4_, 6.0 g/L yeast extract and 0.1 g/L ampicillin) was grown at 30 °C to an OD_600_ of 5.0 and used for inoculation. The cells were cultured in a 10 L jar fermenter (Takasugi Seisakusho Co. Ltd., Tokyo, Japan) containing 5.7 L of medium (12.0 g/L glucose, 9.0 g/L KH_2_PO_4_, 15 g/L yeast extract, 0.04 g/L FeSO_4_ · 7H_2_O, 0.04 MnSO_4_ · 5H_2_O, 0.04 CaCl_2_ · 2H_2_O, GD-113 (Nof Corporation, Japan)) at 37 °C. The initial OD_600_ was 0.05. After the initial glucose was consumed, a feeding solution containing 65.9 w/v% glucose was pumped into the fermenter at a feeding rate of 50 mL/h. The dissolved oxygen concentration was kept at 20% air saturation. The pH was kept at pH 6.9. After 24 h, protein expression was induced by adding isopropyl-β-D-thiogalactoside (IPTG) to a final concentration of 0.1 mM. The cells were harvested by centrifugation at 11,000 x g for 15 min at 24 h after induction. For the recombinant spidroins, the harvested cells were washed out with 20 mM Tris-HCl, pH 7.4, and centrifuged at 11,000 x g, and the supernatant was discarded. The cells were resuspended in 20 mM Tris-HCl, pH 7.4, 100 mM NaCl, pH 8.0 with 0.9 µg of DNase per wet cell weight (g), 82 µg of lysozyme per wet cell weight (g) and 0.2 mL of phenylmethylsulfonyl fluoride (PMSF) per wet cell weight (g). Cells were agitated overnight at 37 °C. The cells were resuspended in buffer solution (50 mM Tris-HCl, 100 mM NaCl, pH 8.0) with 3 w/v% sodium dodecyl sulfate (SDS), and the cell suspension was centrifuged at 11,000 x g for 30 min at room temperature. The pellets were dissolved in DMSO containing 1 M LiCl at 60 °C for 30 min. The solution was centrifuged at 11,000 x g for 30 min at room temperature. Ethanol precipitation was performed at room temperature. The mixture was centrifuged at 11,000 x g for 10 min. The precipitate was washed with reverse-osmosis purified (RO) water three times, followed by lyophilization. For SpiCE-NMa1, the harvested cells were suspended in sodium phosphate buffer (50 mM sodium phosphate, 300 mM NaCl, pH 7) with 7.5 M urea, PMSF and DTT and then lysed by a high-pressure homogenizer (GEA Niro Soavi), followed by centrifugation. The supernatant was filtered through a 0.44 µm filter and loaded onto a column packed with Nuvia IMAC (Bio-Rad). After loading, the resin was washed with 100 mM sodium phosphate buffer with 7.5 M urea, DTT, Triton-X100 and 15 mM imidazole and then eluted using a linear gradient of imidazole concentrations from 15 mM to 500 mM. The eluents were collected and dialyzed against buffer A (20 mM Tris-HCl, 7.5 M urea, 50 mM NaCl, DTT and Triton-X100, pH 7.4). The dialyzed sample was loaded onto a 5 mL Bio-Scale Macro-Prep High Q Column (Bio-Rad). The resin was washed with 20 mM Tris-HCl buffer containing 7.5 M urea, 80 mM NaCl, 1 mM DTT and Triton-X100, pH 7.4, and then eluted using a linear gradient of NaCl concentration from 80 mM to 1 M. The eluents were collected and loaded onto Nuvia IMAC resin again and eluted using a linear gradient of imidazole concentration from 15 mM to 500 mM. The eluents were dialyzed against RO water and then lyophilized.

### Artificial recombinant films

HIFP was used as the solvent for the film synthesis. Recombinant MaSps were mixed in 8 mL HFIP to a total of 100 mg and then dried at room temperature for 16 h in a dish (Fig. S13). Composite films containing SpiCE-NMa1 were produced by mixing the MaSp2 solution with recombinant SpiCE-NMa1 at 1, 3, or 5 wt% each.

### Artificial recombinant silks

Recombinant MaSp2 protein of 22 wt% (1.66 g) was dissolved in 5.89 g DMSO (4 wt% LiCl) and incubated at 90 °C for 8 h. For composite silk containing SpiCE-NMa1, recombinant MaSp2 protein of 22 wt% and recombinant SpiCE-NMa1 of 0.012 g were dissolved in 4.2 g DMSO. The dope was aspirated with a N_2_ pump and D=0.1 mm needle at 70 °C and spun directly into baths containing MeOH/DMSO 30 v/v% (5 °C), MeOH (25 °C), and water (25 °C). Only MaSp2/SpiCE-NMa1 composite silk could be produced because of the low yield of recombinant MaSp1 and MaSp3 proteins.

### Wide-angle X-ray scattering (WAXS) measurement

The crystalline samples were characterized by a synchrotron WAXS measurement at the BL45XU beamline of SPring-8, Harima, Japan (75). The X-ray energy was set at 12.4 keV (wavelength: 0.1 nm), and the distance between the sample and the detector was approximately 257 mm. For the diffraction patterns, an exposure time of 10 sec was used. Using Fit2D software (76), the obtained data were converted into one-dimensional radial integration profiles and corrected by subtracting the background scattering. The degree of crystallinity was calculated from the area of the crystal peaks divided by the combined area of the crystal peaks and the amorphous halo by fitting the Gaussian function using Igor Pro 6.3.

### Measurement of mechanical properties

The tensile properties of the fibers were measured using an EZ-LX universal tester (Shimadzu, Kyoto, Japan) with a 1 N load cell at a strain rate of 10 mm/min (0.033 s^-1^) at 25 °C and 48% relative humidity. For each tensile test, the cross-sectional area of an adjacent section of the fiber was calculated based on the SEM images. The transparency of the film was measured at selected wavelengths between 350 and 600 nm using a UV instrument. Each fiber was attached to a rectangular piece of cardboard with a 5 mm aperture using 95% cyanoacrylate.

Tensile tests of the recombinant silk fibers were performed using a mechanical testing apparatus (Automatic Single-Fiber Test System FAVIMAT+, Textechno H. Stein GmbH & Co. KG, Mönchengladbach, Germany) with a 210 cN load cell at a strain rate of 10 mm/min (0.016 s^-1^) at 20 °C and RH 65%. From each fiber sample, diameters were measured at different sites (n = 6) by microscopic observation (Nikon Eclipse LV100ND, lens 150×0) to calculate the cross-sectional area of an adjacent section of the fibers.

## Acknowledgments

The authors thank Akio Tanikawa for morphological identification of spiders and helpful comments about phylogenetic discussion and thank the Malagasy Institute for the Conservation of Tropical Environments (MICET), the Ministry of Environment and Sustainable Development of Madagascar, Mention Zoologie et Biologie Animale (MZBA), Université d’Antananarivo, and Madagascar National Parks (MNP) for their cooperation and permission to conduct sampling in Madagascar. The authors also thank Yuki Takai, Nozomi Abe, Yuki Onozawa, and Naoko Ishii for technical support in the sequencing and proteome analysis, Hongfang Chi, Haruka Funayama, Kaori Yaosaka, Ryota Sato, and Hironori Yamamoto for technical support in the fiber spinning and analysis of fibers, Hiroshi Kano, Ryoko Sato, and Hideki Nishijima for the development of reeling machines, and Hiroyasu Masunaga for his technical support in SPring-8 measurements. H.N., M.T., K.N., and A.K. are supported by the ImPACT Program of the Council for Science, Technology and Innovation (Cabinet Office, Government of Japan). N.K. is supported by a Nakatsuji Foresight Foundation Research Grant, the Sumitomo Foundation (190426), The Uehara Memorial Foundation, Grant-in-Aid for Scientific Research (B). N.K., M.M., M.T., and A.K. are supported by research funds from the Yamagata Prefectural Government and Tsuruoka City, Japan. Y.Y. is supported by Japan Society for the Promotion of Science (JSPS) KAKENHI Grant Number JP18J21155, and K.N. is financially supported by a Grant-in-Aid for Transformative Research Areas (B).

## Supplementary Information

### Supplementary Information Text

#### Improvement of the *T. clavipes* (NepCla1.0) genome and spidroins

##### Genome improvement

The first *T. clavipes* genome was reported in (1). This paper made a significant contribution to spidroin research, with the spidroin catalog of orb-weaving spiders constructed from the draft genome demonstrating that spidroins constitute a complex gene family. However, the sequencing of large spider genomes is very challenging, and classic technologies alone can obtain only limited information. In addition, the risk of contamination must always be considered in genome sequencing. Steinegger and Salzberg conducted a comprehensive analysis of public databases and reported millions of contaminants in genome sequences (2). In that report, they described that all 504 contigs (over 24 M residues) in the *T. clavipes* genome (NepCla1.0) were identified as possible contaminants (2). We also submitted the *T. clavipes* genome (NepCla1.0) to BlobTools analysis (3) to detect possible contaminants and showed that 4,935 scaffolds (over 44 Mb), including the top two long scaffolds (MWRG01000001.1 and MWRG01000002.1 and, total 2.89 Mbp), may be of bacterial origin (Fig. S4). In addition to contamination, the completeness of the genome assembly and the predicted gene set must be carefully evaluated. N50 and BUSCO are de facto benchmarks for evaluating the completeness and credibility of genome assembly or gene sets and should always be evaluated. The BUSCO completeness of the *T. clavipes* genome (NepCla1.0: GCA_002102615.1_NepCla1.0) and *T. clavipes* gene set (NepCla1.0: 50,846 genes published as MWRG01_proteins_060117) were 67.4% and 62.2%, respectively, for the Arthropoda lineage; they were 72.1% and 60.4%, respectively, even for the eukaryote lineage. The N50 and N90 scaffold sizes were 62,959 bp (#9,532) and 5,317 (#65,231), respectively.

The *T. clavipes* genome we report here is the result of the hybrid assembly using newly sequenced reads with short-read sequencing to guarantee accuracy and a long-read sequencing to guarantee structure. The removal of contaminants was strictly conducted using BlobTools based on homology searches (Fig. S2). We then presented in this paper a sufficiently improved genome with an N50 of 738,925 bp (#1,112) and BUSCO of 93.0%. The BUSCO score of the predicted gene set was 88.2%, which is considered to be high enough quality to be used. Spidroin improvement

All known *T. clavipes* spidroins have been isolated based on short-read data obtained from gDNA or cDNA sequencing. The typical spidroin structure is one in which repeat domains are sandwiched between the nonrepeated terminal domains, and a reasonable spidroin ortholog group lineage has been shown. However, full-length spidroin genes have not been sequenced directly, and classic assemblies from short reads alone may have collapsed the repeat domains. We confirmed all spidroin lengths based on the long reads obtained by Nanopore direct sequencing and validated the full-length structures. Accordingly, a new proposal for nomenclature is now possible. The spidroin gene set (NepCla1.0) was systematized, and the spidroin genes were also given appropriate names. In particular, our comparative and phylogenetic analysis showed that Sp-907 and Sp-74867 are MaSp family 3 spidroins.

##### Spidroin nomenclature

According to N-terminal domain similarity, we defined the following nomenclature system for spidroins in this paper. Names of individual spidroin members are represented as “SPnXm”, where SP is the root name (MaSp, MiSp, AgSp, PySp, Flag, CySp, or AcSp), n is an integer representing a family, X is a single letter denoting a subfamily, and m is an integer representing an individual family member (isoform) (Fig. 1 and Fig. S5).

There have been many previous reports of the existence of multiple MaSp families. In addition to the classic MaSp families 1 and 2, a recent high-throughput sequencing approach has found a third family (family 3) in *A. ventricosus*, *Araneus diadematus*, *Argiope argentata*, *Latrodectus hesperus*, and *Parasteatoda tepidariorum* (4, 5) and MaSps a-h in *T. clavipes* (1). Furthermore, families 4 and 5 have also been reported in *Caerostris darwini* with unique repeat motifs (6). These sequences were also sorted into appropriate families by our comparative analysis (Fig. S7).

For example, from previous studies, MaSp-abcdefgh and MiSp-abcd had been reported in alphabetical order, and Sp-907 (PRD35000.1) and Sp-74867 (PRD19552.1) had been reported as just spidroin proteins from the previous *T. clavipes* genome (NepCla1.0). Here, all of these proteins except for the very short MaSp-e were classified based on the phylogenetic analysis (Fig. S7), as follows:

MaSp-b/c (PRD18716.1/PRD18936.1), MaSp-a (PRD23950.1), MaSp-d (PRD23750.1), MaSp-h (PRD20448.1), MaSp-g (PRD24320.1), MaSp-f (PRD27696.1), MiSp-b/c/d (PRD30268.1/PRD24510.1/PRD18914.1), MiSp-a (PRD23654.1), Sp-907 (PRD35000.1), and Sp-74867 (PRD19552.1) are MaSp1A3, MaSp1C, MaSp1B1, MaSp2A2, MaSp2A3, MaSp2B1, MiSp1A, MiSp1B, MaSp3C1, and MaSp3B1, respectively.

---

**Fig. S1.**
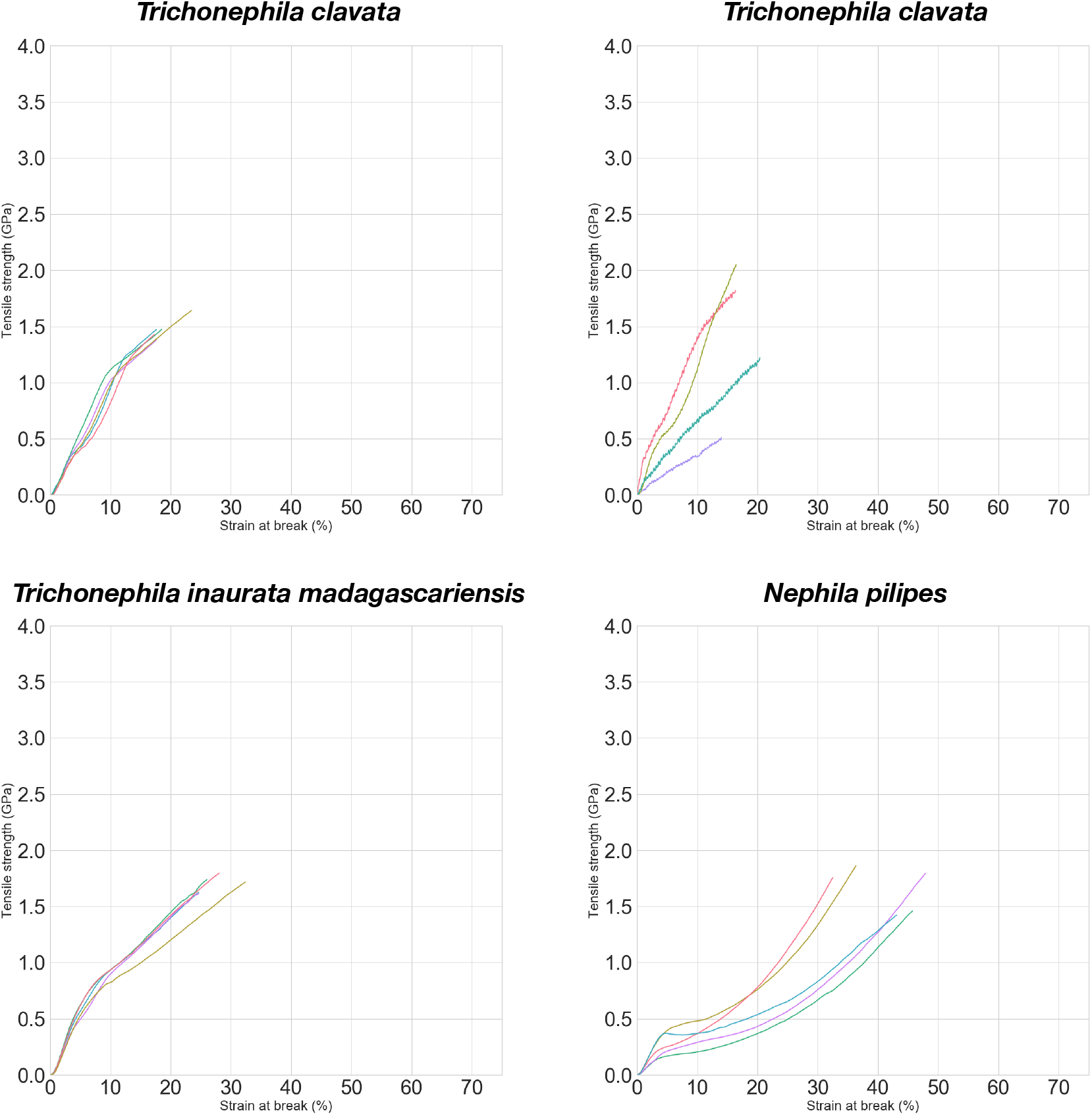
Stress-strain curves of dragline silk in four golden silk orb-weavers (subfamily Nephilinae). The results of tensile tests of dragline silks from *Trichonephila clavata* (toughness: 169 ± 39 MJ/m^3^; Young’s modulus 11.80 ± 0.82 GPa; tensile strength 1.51 ± 0.10 GPa; strain at break: 19.2 ± 3.3%), *Trichonephila clavipes* (toughness: 131 ± 36 MJ/m^3^; Young’s modulus: 10.40 ± 2.86 GPa; tensile strength: 1.56 ± 0.24 GPa; strain at break: 17.1 ± 3.3%), *Trichonephila inaurata madagascariensis* (toughness: 285 ± 41 MJ/m^3^; Young’s modulus: 14.60 ± 1.30 GPa; tensile strength: 1.73 ± 0.07 GPa; strain at break: 27.2 ± 3.2%), and *Nephila pilipes* (toughness: 292 ± 40 MJ/m^3^; Young’s modulus: 7.00 ± 2.50 GPa; tensile strength: 1.72 ± 0.34 GPa; strain at break: 41.8 ± 6.8%).

**Fig. S2.**
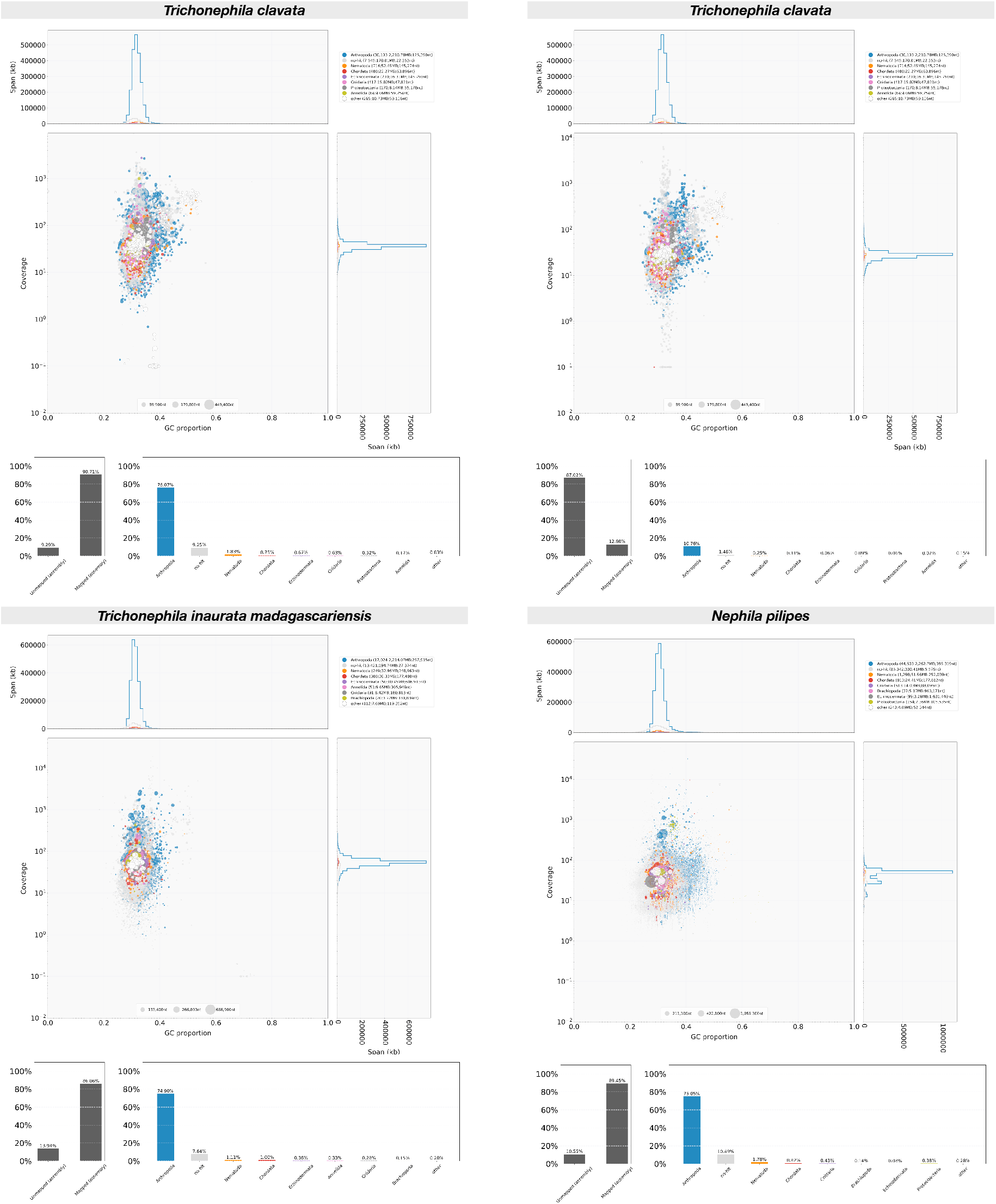
Graphical representation of the contaminant analyses. Results from the contamination check of the *T. clavata*, *T. clavipes*, *T. inaurata madagascariensis*, and *N. pilipes* genome assemblies using BlobTools (3) and Diamond (7). The BlobTools plot shows the taxonomic affiliation at the phylum rank level, distributed according to GC% and coverage.

**Fig. S3.**
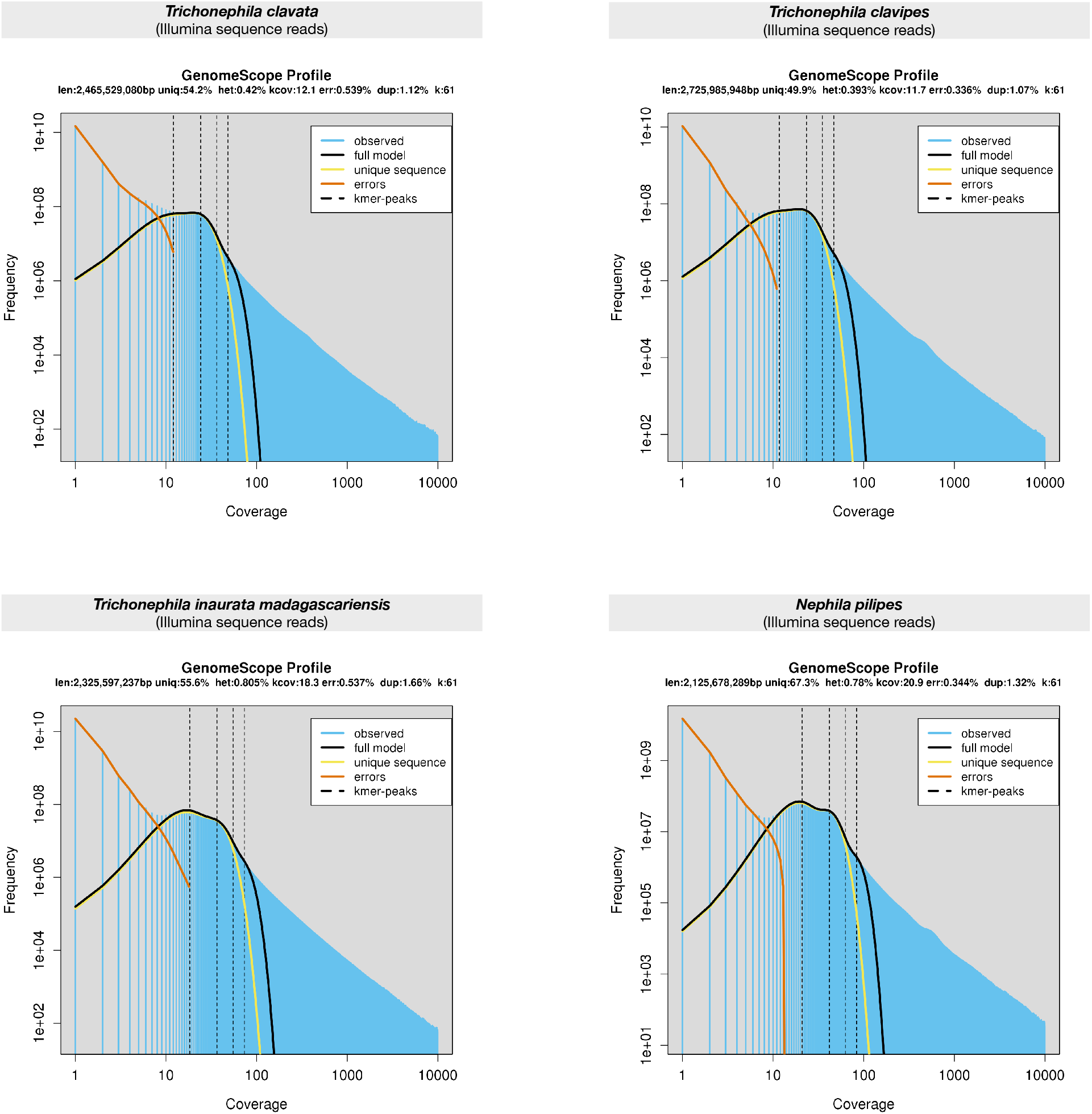
Graphical results of genome size estimation. The size of each genome was estimated by GenomeScope (8), providing a k-mer analysis (k = 61) based on raw Illumina sequence reads obtained from genome sequencing.

**Fig. S4.**
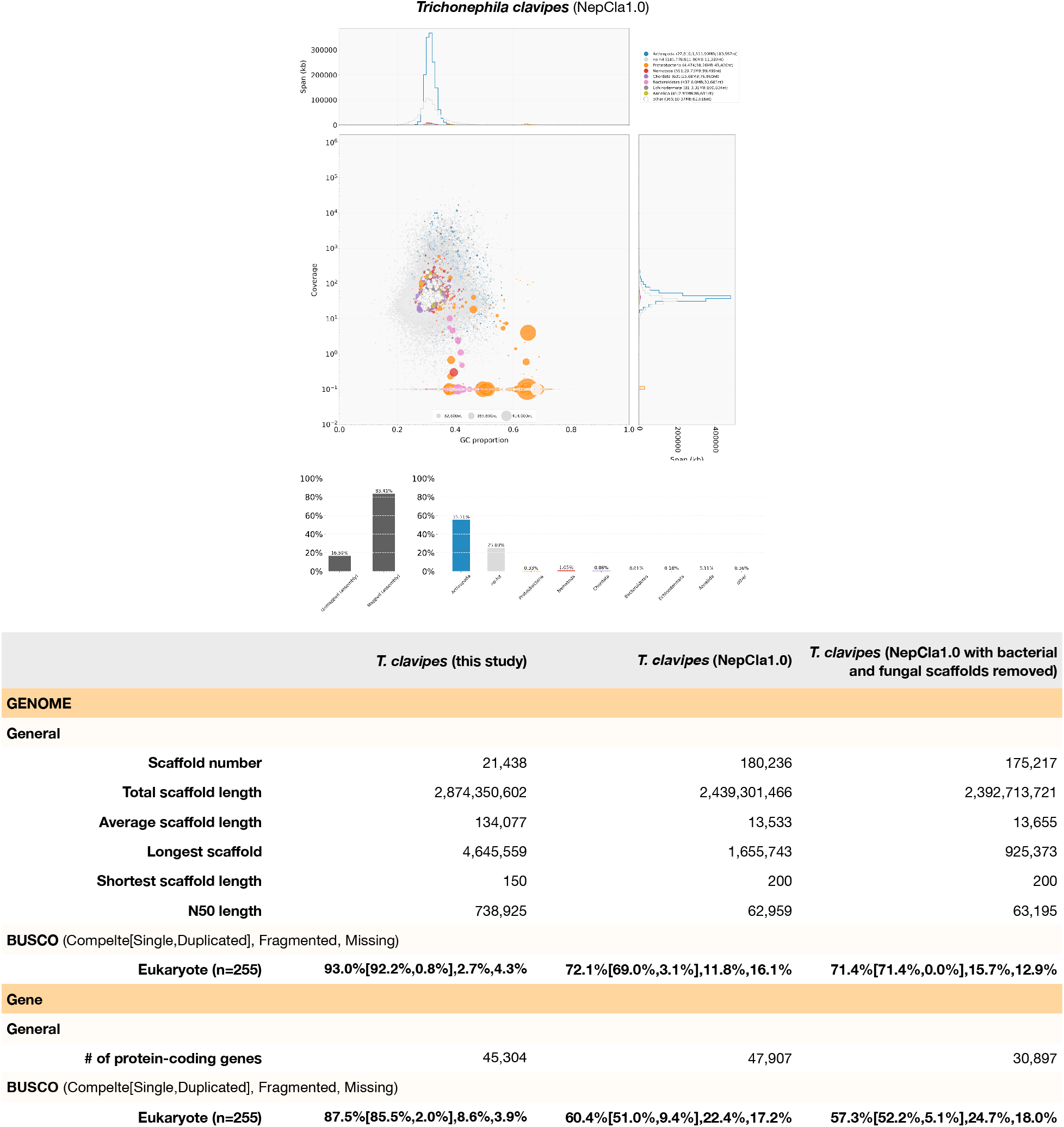
Comparison of *T. clavipes* genome statistics from this and a previous study. The panel shows the results of contamination detection in the previously reported *T. clavipes* draft genome (NepCla1.0), and the table shows the genome statistics. The large orange circles and many pink circles indicate the scaffolds classified as Proteobacteria (38.26 Mb) and Bacteroidetes (6.0 Mb), respectively. The bottom table shows the genome statistics of our *T. clavipes* draft genome (“this study”, Table 1) and the previous *T. clavipes* draft genome with and without contamination scaffolds (“NepCla1.0” and “NepCla1.0 with bacterial and fungal scaffolds removed”).

**Fig. S5.**
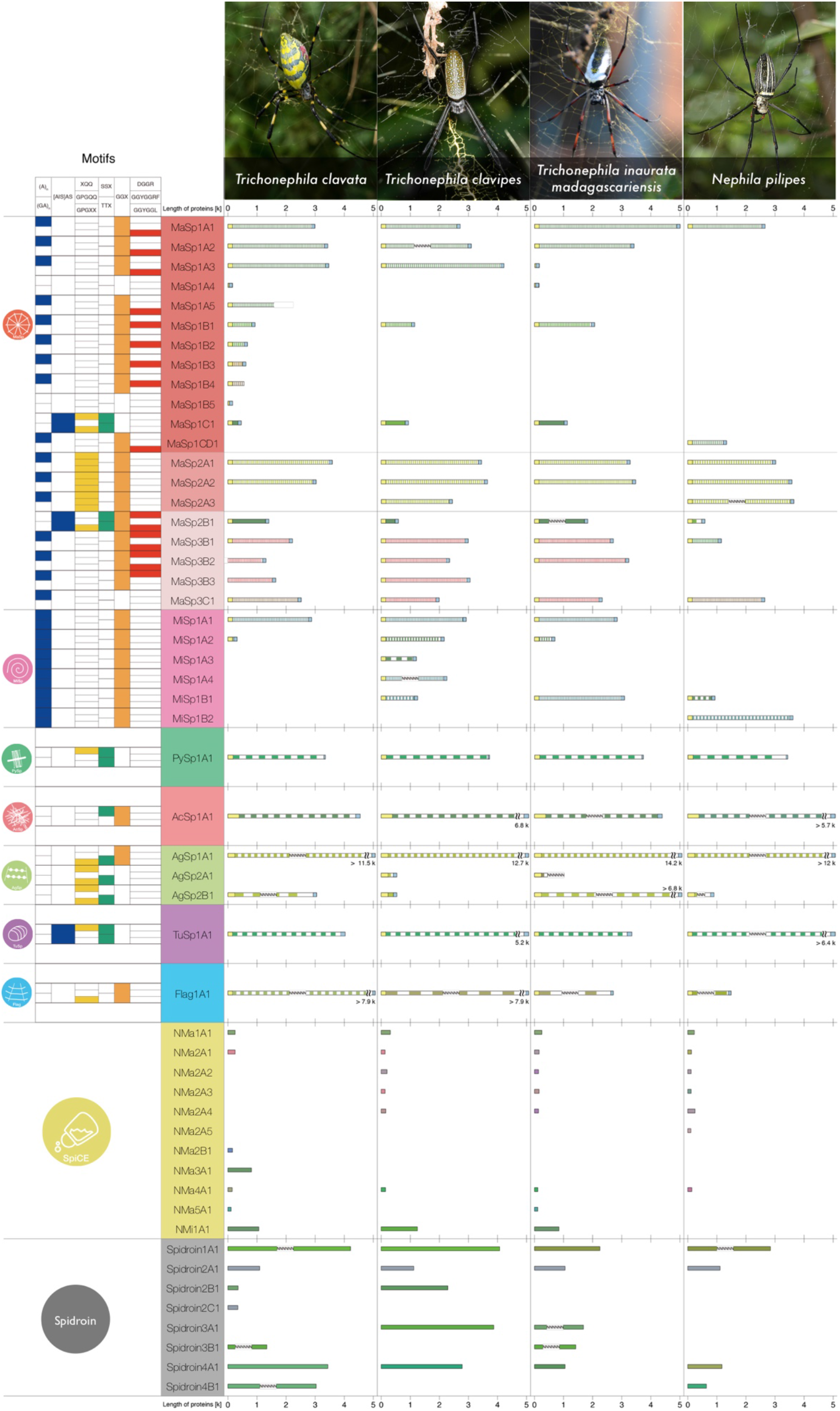
Spidroin and SpiCE catalog. This panel shows the spidroin and SpiCE sequence characteristics and structures obtained from four Nephilinae draft genomes. The icons in the first column represent spidroin groups. The second column represents the motif variety in the repeat domains: β-sheet (An, (GA)n, and AS), blue; β-turn (XQQ, GPGQQ, and GPGXX), yellow; spacer (SSX and TTX), green; 310 helix (GGX), orange; and MaSp motif (DGGR, GGYGGRF, and GGYGGL), red. The spidroin structure columns show the N/C-terminal (yellow and blue box) and repeat domains, and each structure is drawn to scale. The number, width, and color of stripes reflect the number, size, and motif of repeats. Assembly gaps are indicated as “N”.

**Fig. S6.**
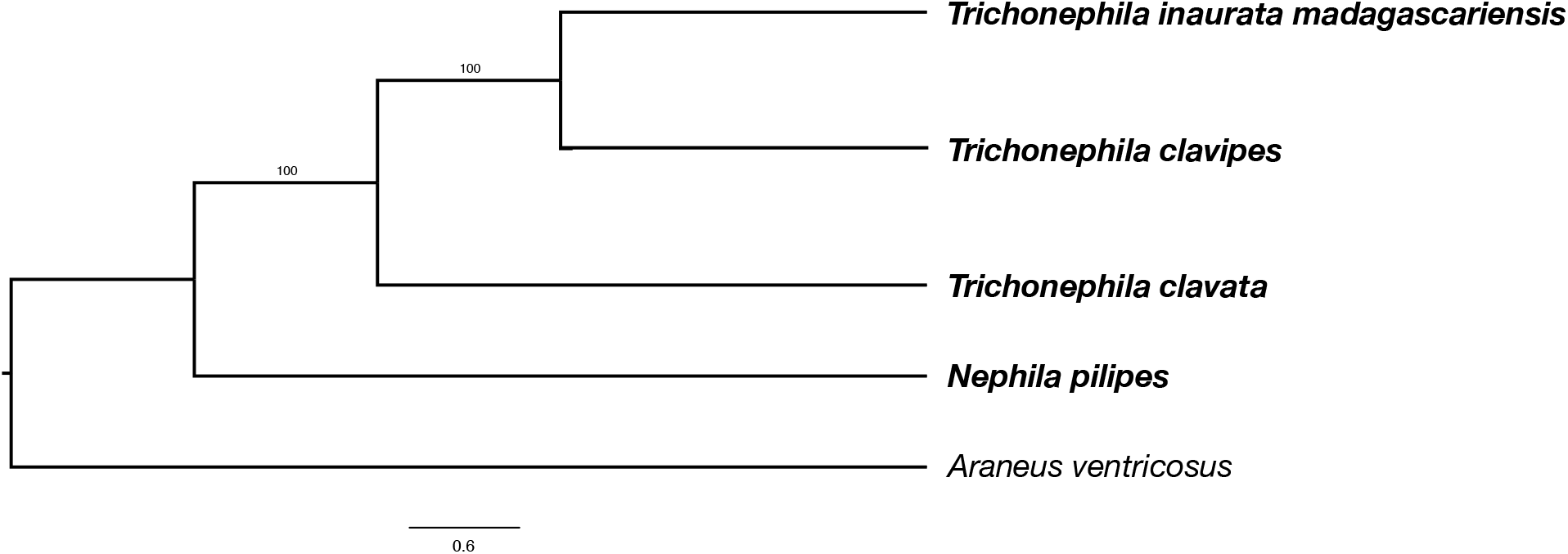
Phylogenetic tree of four sequenced Nephilinae species. Phylogenetic tree based on a protein sequence corresponding to the 1,031 core orthologous gene set in arthropod (9).

**Fig. S7.**
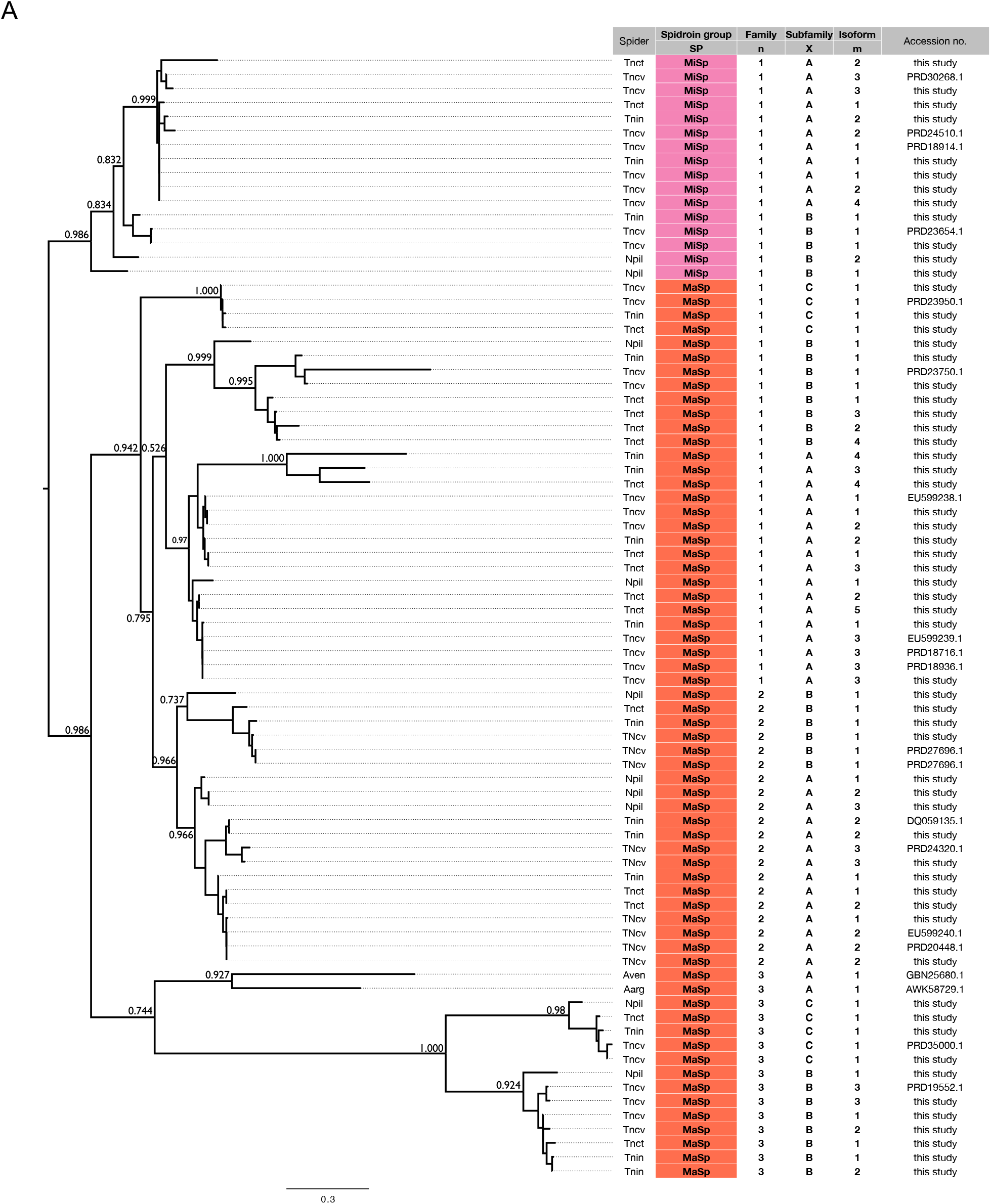
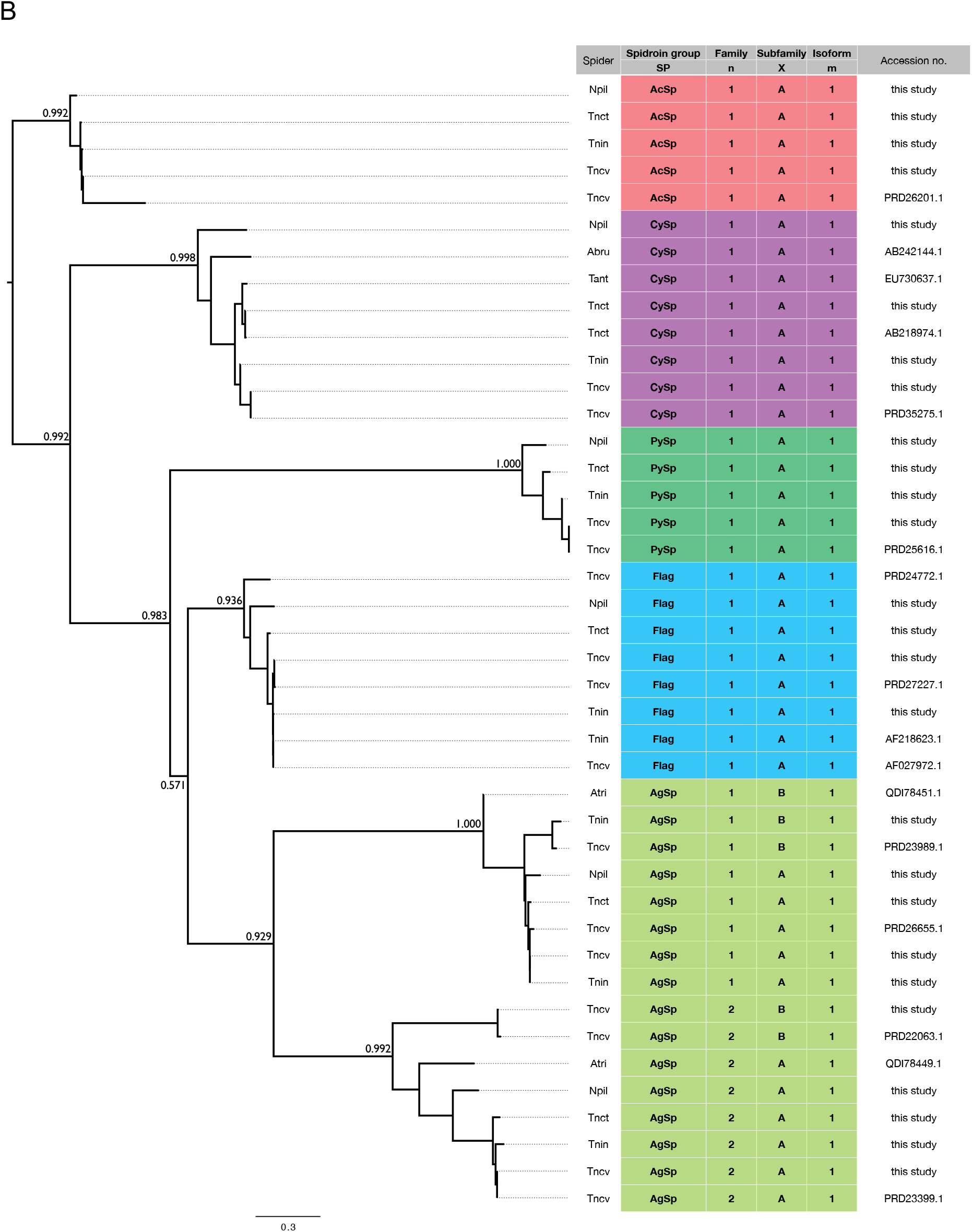
Systemic relationships underlying new nomenclature. These phylogenetic trees are constructed on the basis of the 150 aa N-terminal regions of **(A)** the ampullate spidroins (MaSp and MiSp) and **(B)** the other spidroins. The spidroin nomenclature (SPnXm) in this paper is based on this phylogenetic tree. Tnct: *Trichonephila clavata*, Tncv: *Trichonephila clavipes*, Tnin: *Trichonephila inaurata madagascariensis*, Npil: *Nephila pilipes*, Aven: *Araneus ventricosus*, Aarg: *Araneus argentata*, Tant: *Trichonephila antipodiana*, Abru: *Argiope bruennichi*, Atri: *Argiope trifasciata*.

**Fig. S8.**
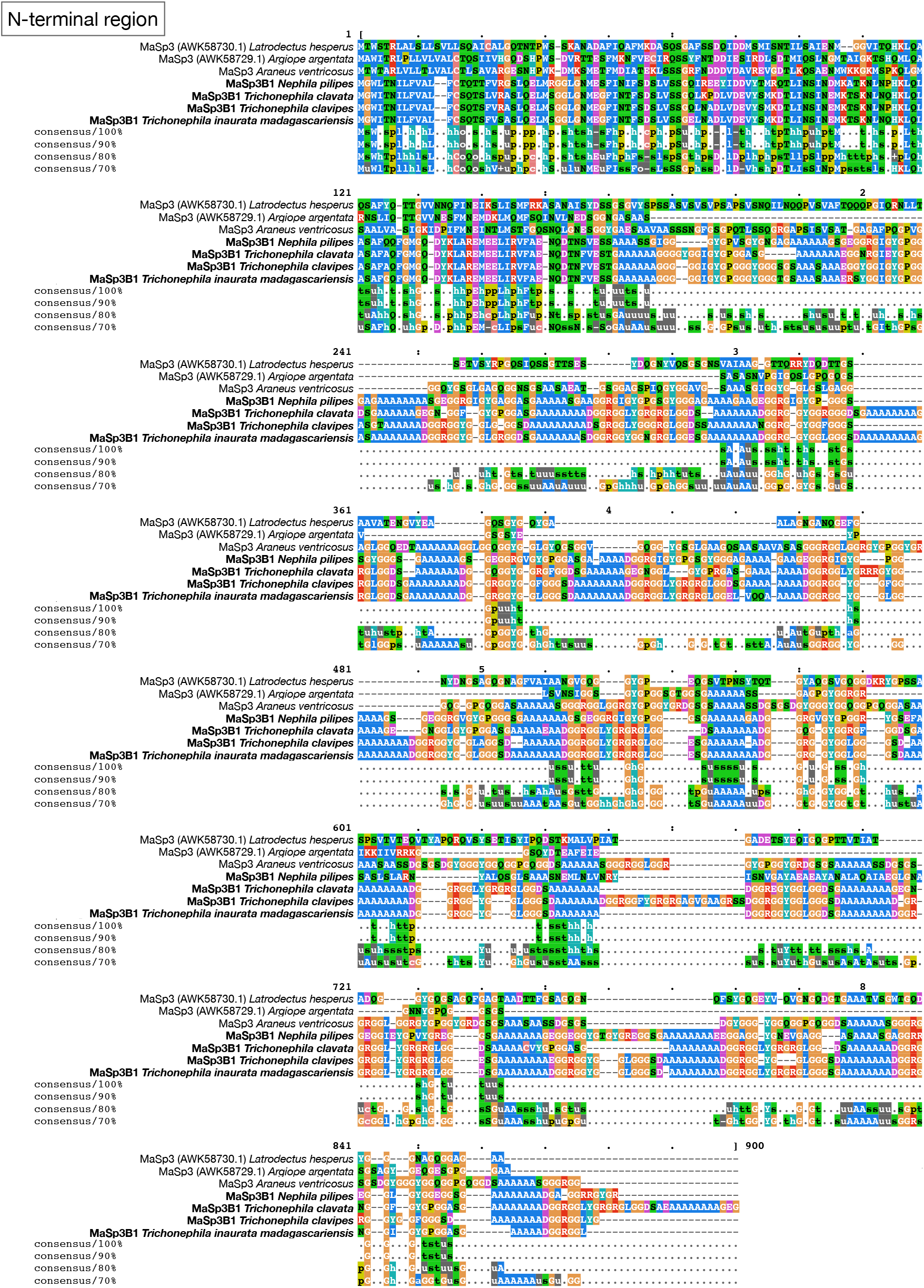
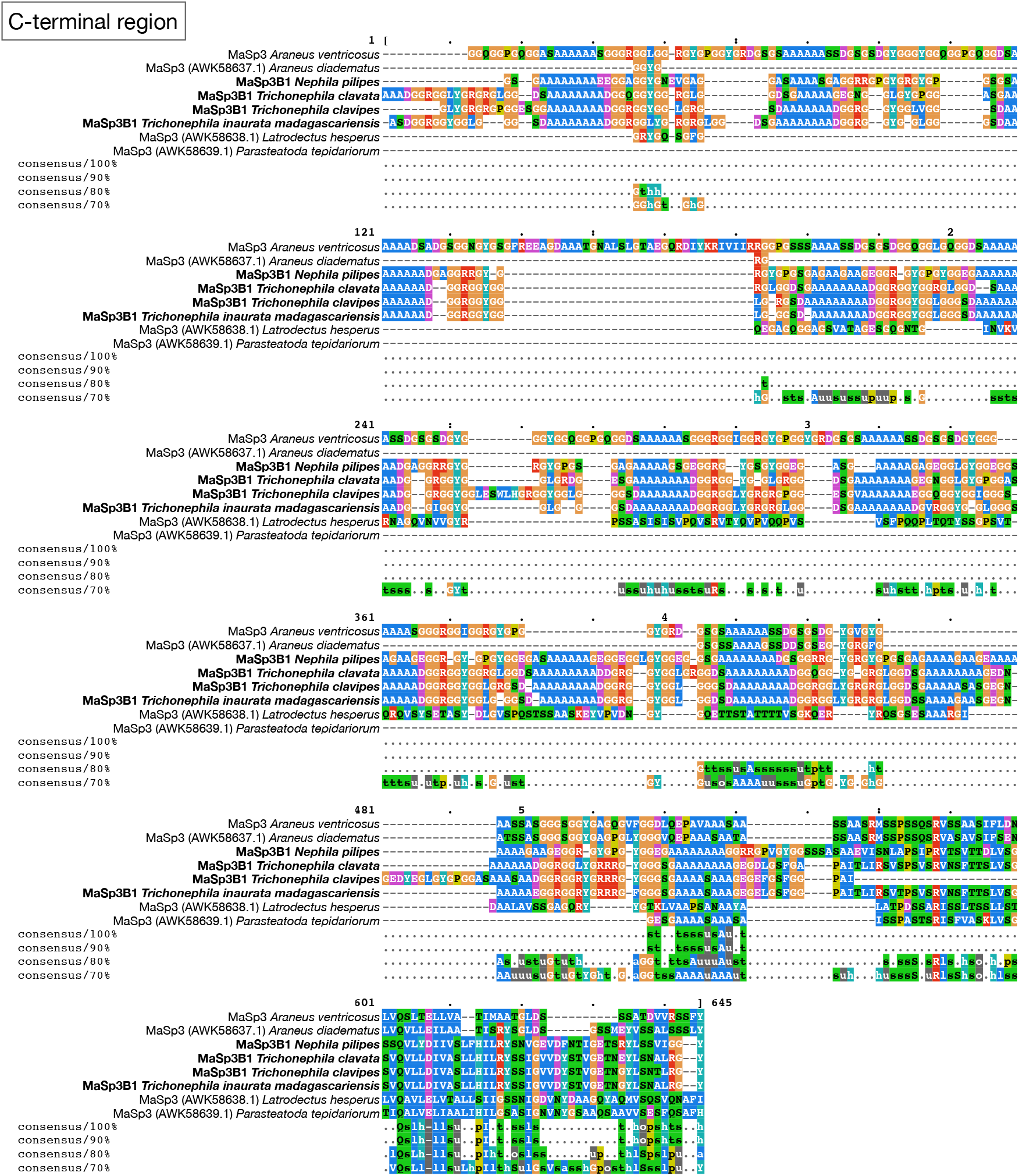
Alignment results of N/C-terminal regions in the MaSp family 3 genes. Two panels show the alignment results of the N- or C-terminal regions of MaSp family 3 genes newly found in subfamily Nephilinae and the known MaSp3 genes. The terminal regions are well conserved and share the same characteristics, with many arginine residues in the repeat domain.

**Fig. S9.**
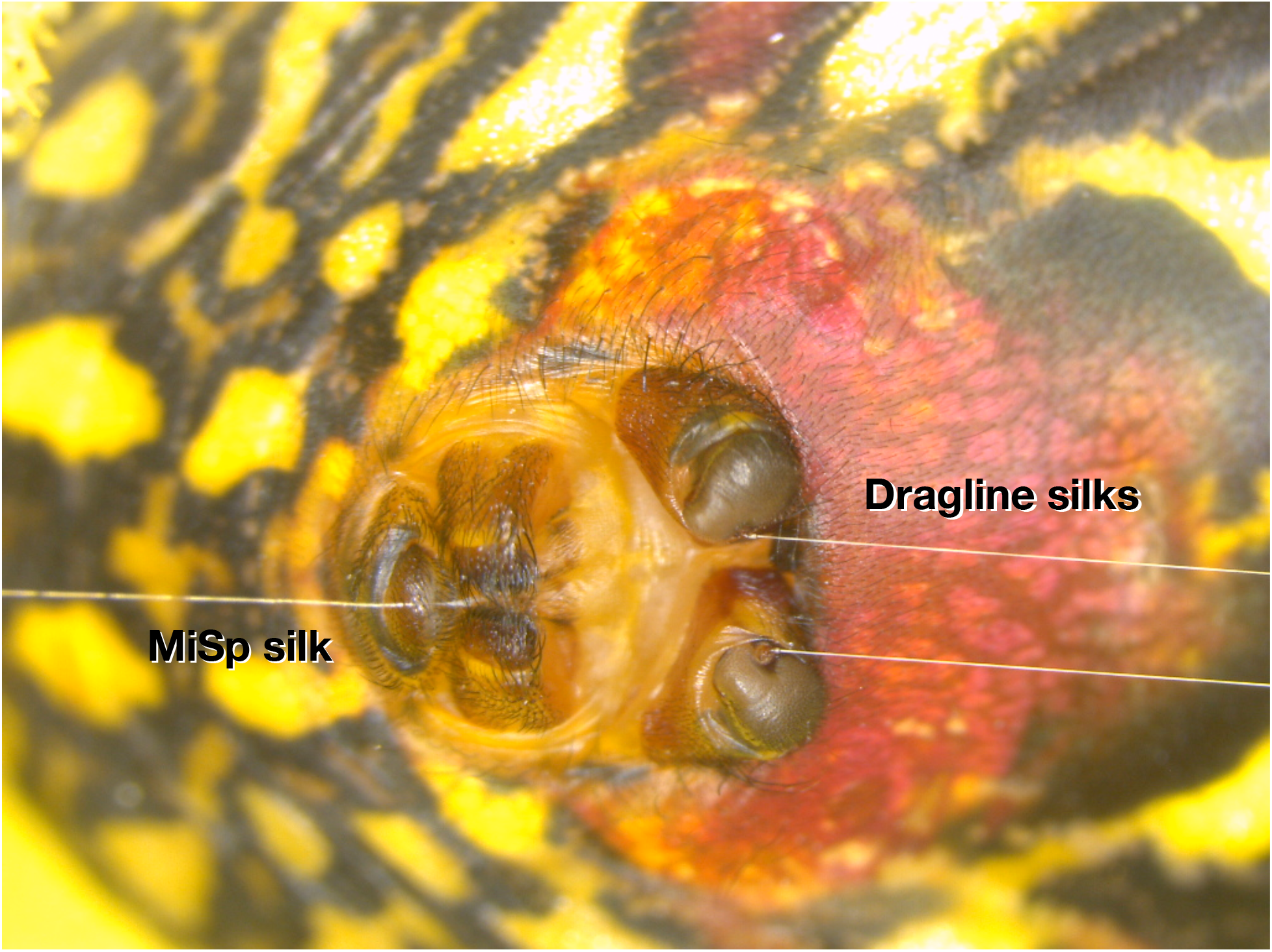
Spinnerets producing dragline (MaSp) and MiSp silk. Dragline silks were reeled directly from the major ampullate gland spigots using a stereomicroscope to avoid mixing with other silks coming out of other spinnerets. This picture is of *T. clavata*.

**Fig. S10.**
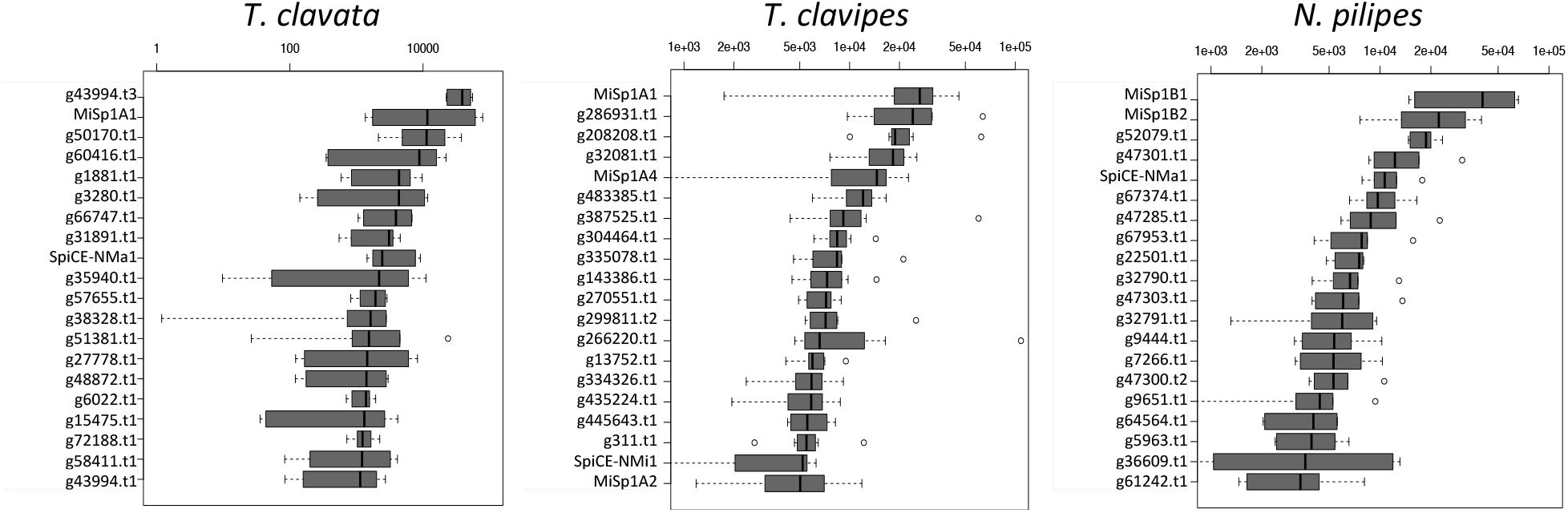
Expression profiling of the minor ampullate silk gland. Expression profiles from the minor ampullate silk gland in *T. clavata* (n = 6), *T. clavipes* (n = 8), and *N. pilipes* (n = 6). The X-axes represent the median of the quantified expression level in TPM (transcripts per million). The y-axes show the top 20 most highly expressed genes in each gland.

**Fig. S11.**
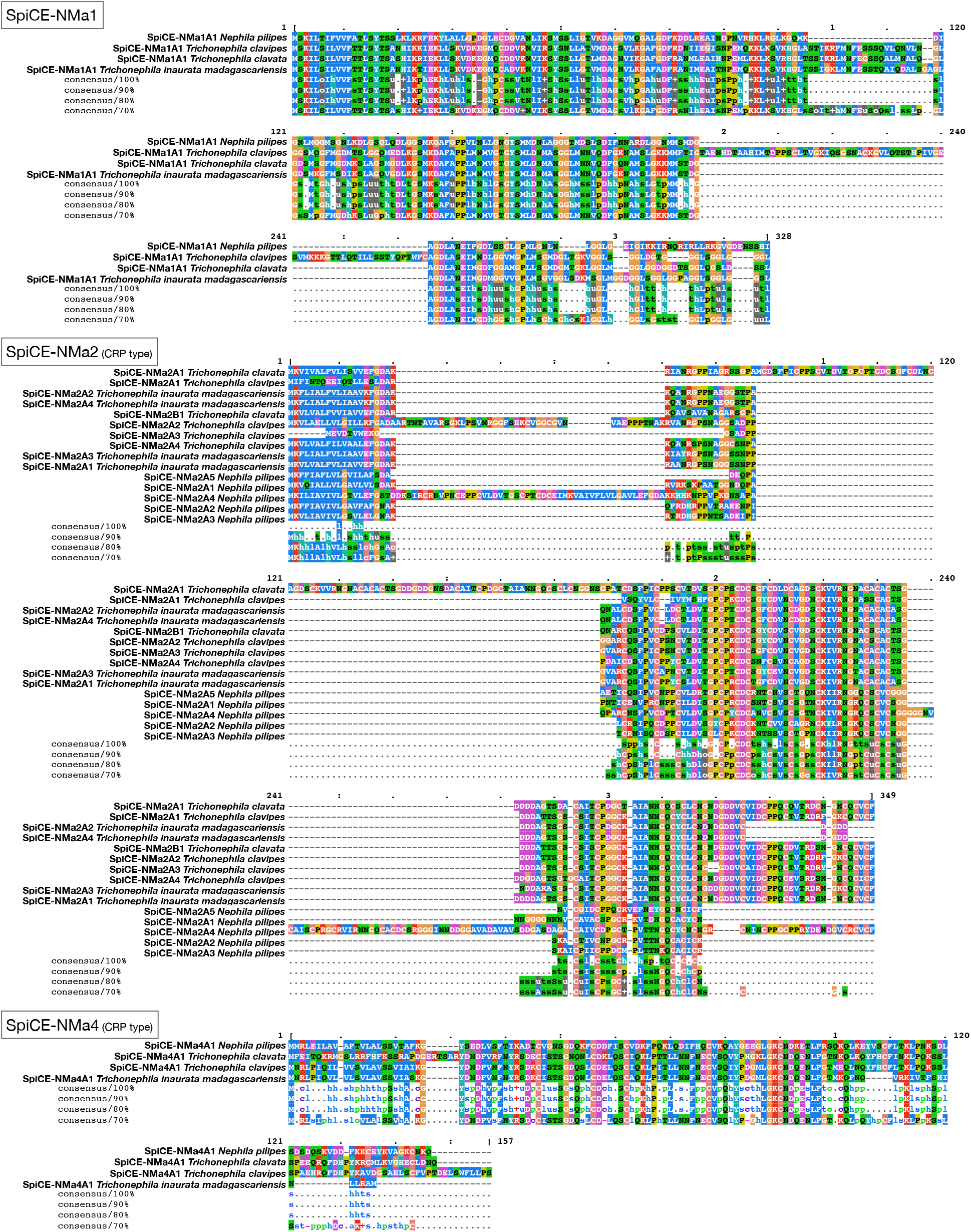
Alignment of SpiCE proteins. These figures show the alignment results of two SpiCE proteins (SpiCE-NMa1, SpiCE-NMa2, and SpiCE-NMa4) in *T. clavata*, *T. clavipes*, *T. inaurata madagascariensis*, and *N. pilipes*.

**Fig. S12.**
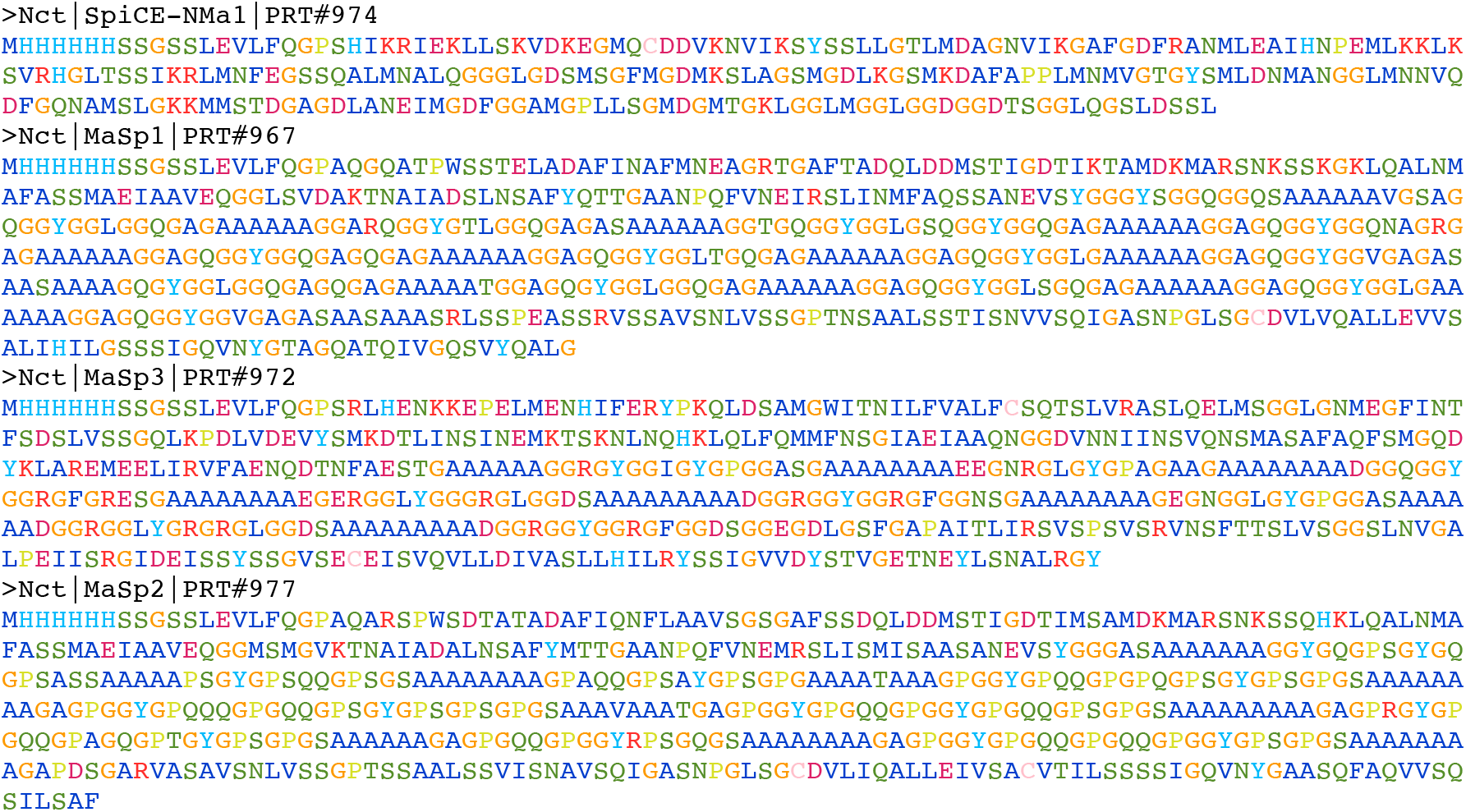
Shortened recombinant amino acid sequences. The optimized amino acid sequences used to produce recombinant proteins (MaSp1, MaSp2, MaSp3, and SpiCE-NMa1). The iteration of the repeat domain in each MaSp gene was modified to be shorter than in the native protein sequences. These genes are composed of a 6x His tag (MHHHHHH), a linker (SSGSS), and an HRV 3C protease recognition site (LEVLFQGP).

**Fig. S13.**
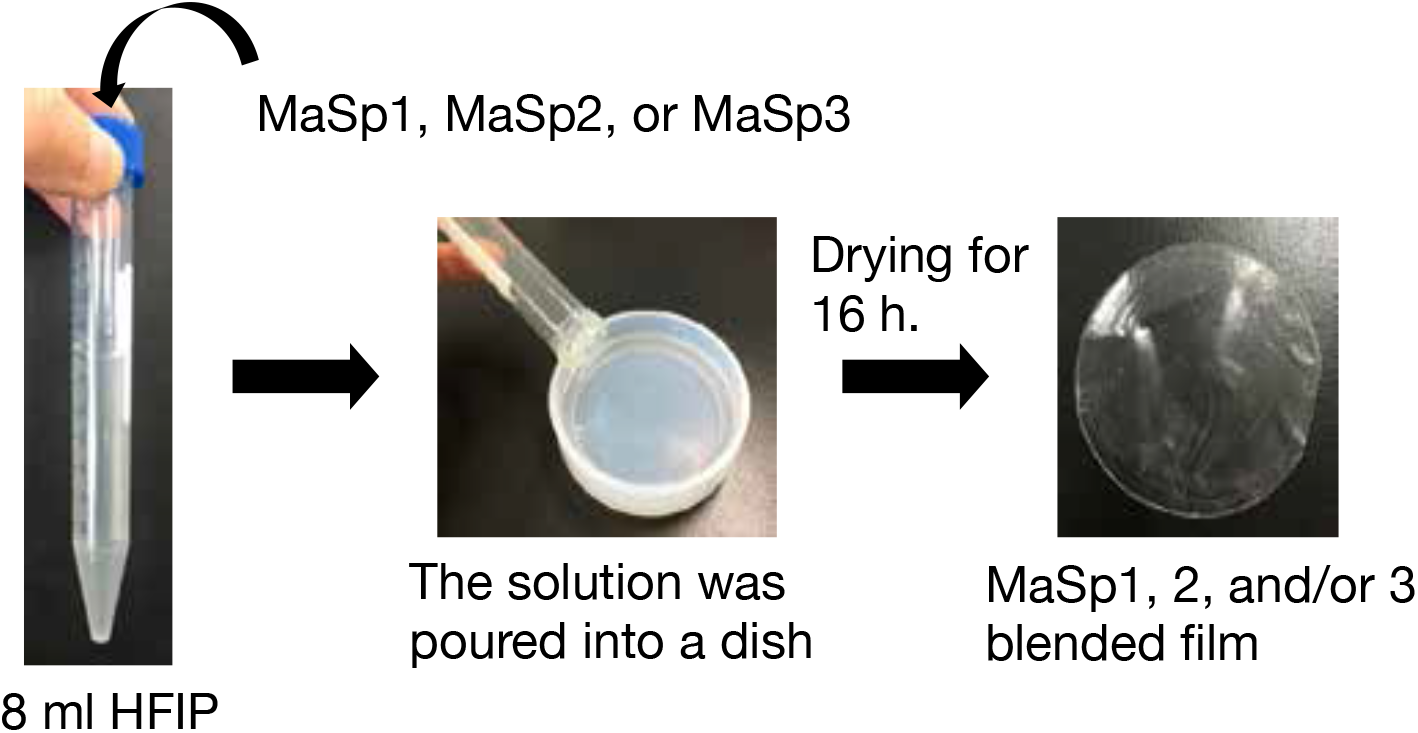
Film formation procedure. This figure represents the composite film production procedure. Recombinant MaSps were mixed in 8 mL HFIP to a total of 100 mg and dried at room temperature for 16 h in a dish. The blending of the recombinant proteins was conducted on the dish.

**Fig. S14.**
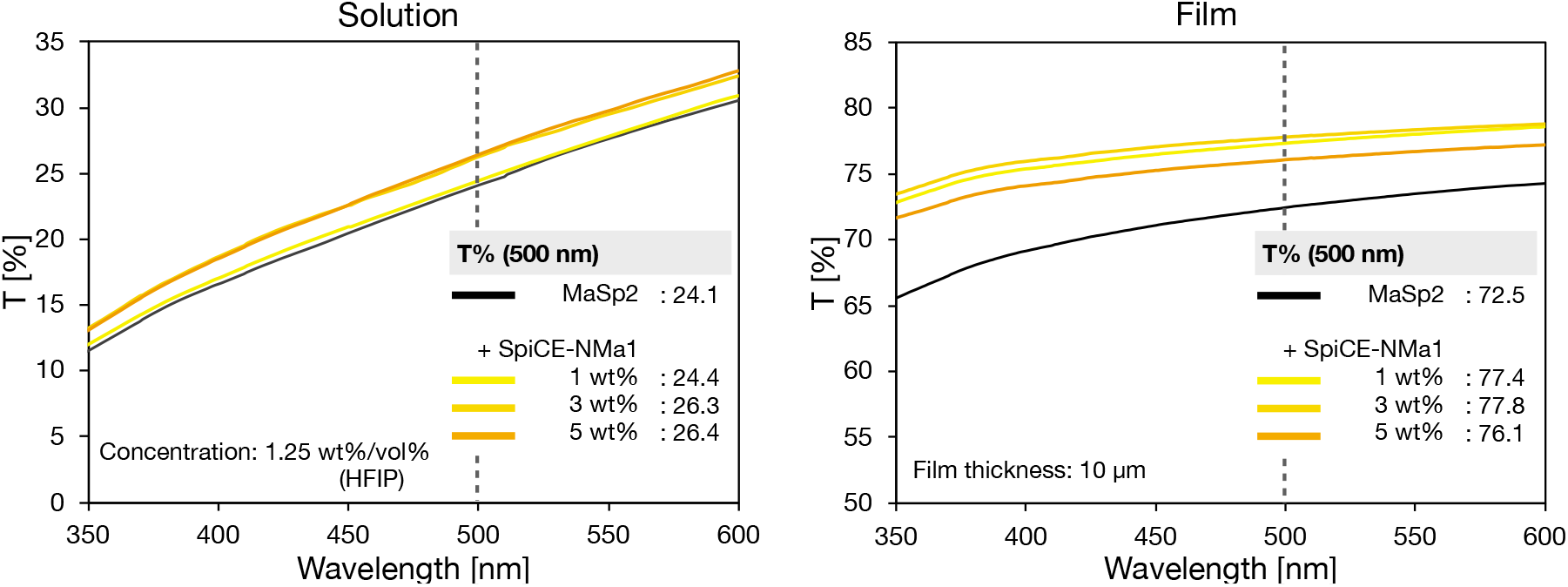
Recombinant protein film transparency. These graphs show the total transparency (%) of MaSp2 and SpiCE composite films with different SpiCE concentrations (0, 1, 3, 5 wt%). The thickness of the used films was 10 µm.

**Fig. S15.**
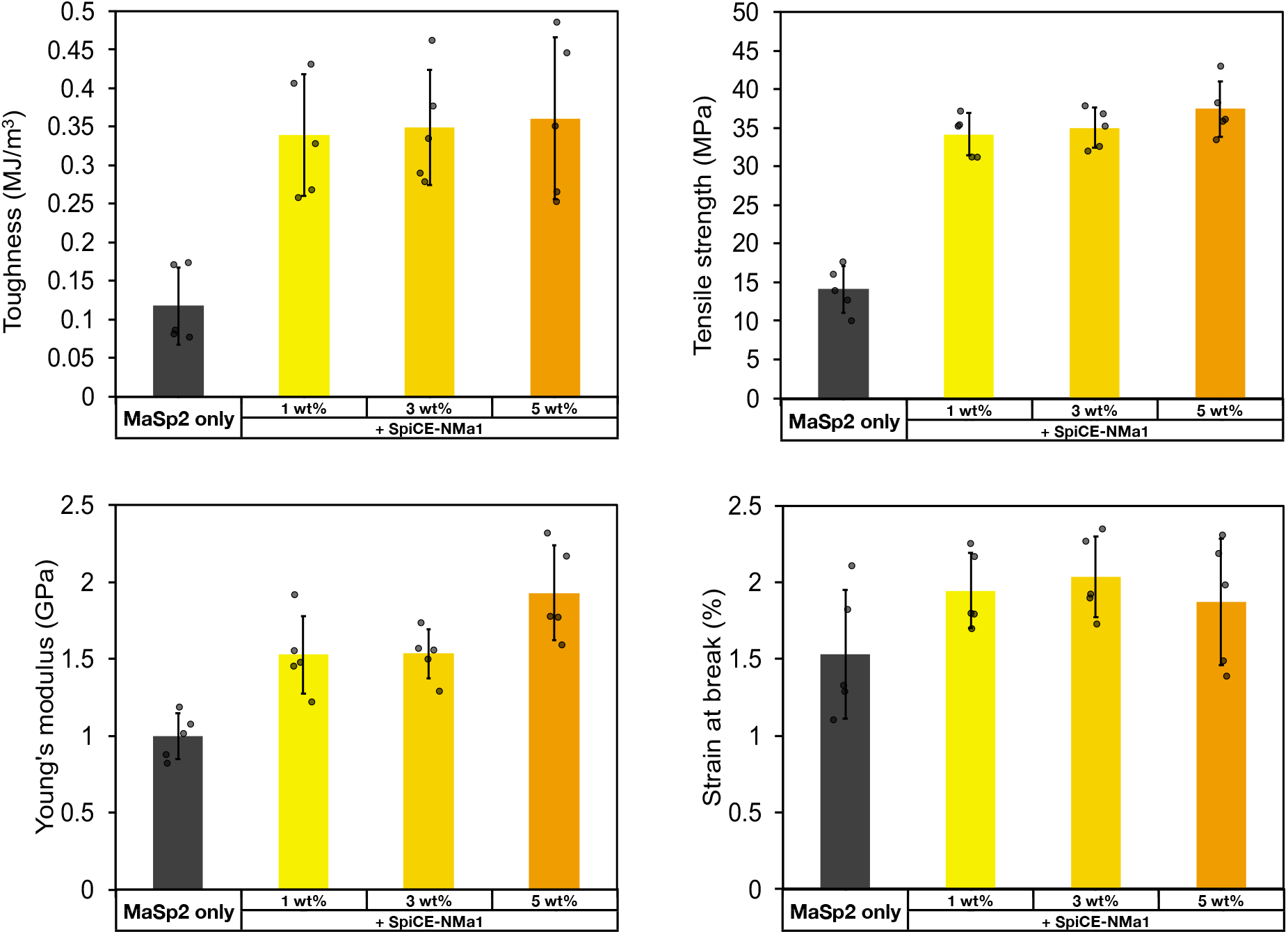
Mechanical properties of recombinant composite films. Graphs showing the tensile properties, such as modulus, strain, strength, and toughness, of MaSp2 and SpiCE protein composite films with different SpiCE concentrations (0, 1, 3, 5 wt%).

**Fig. S16.**
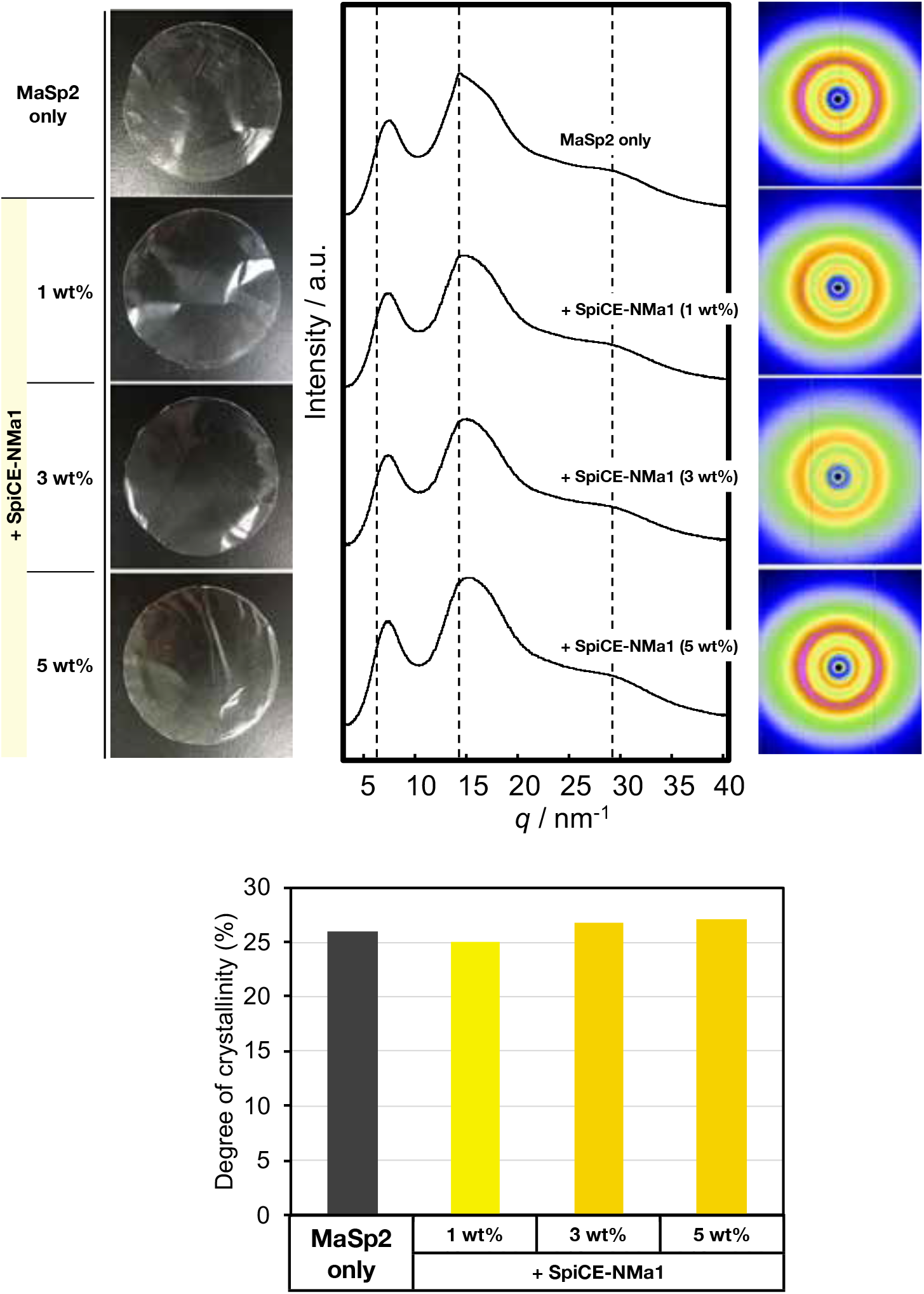
WAXS pattern and crystallinity of recombinant composite films. WAXS measurements of MaSp2 and SpiCE protein composite films under different SpiCE concentrations (0, 1, 3, 5 wt%). The top figure shows one-dimensional radial integration profiles and two-dimensional profiles. The bottom graph shows the degree of crystallinity in each composite film.

**Fig. S17.**
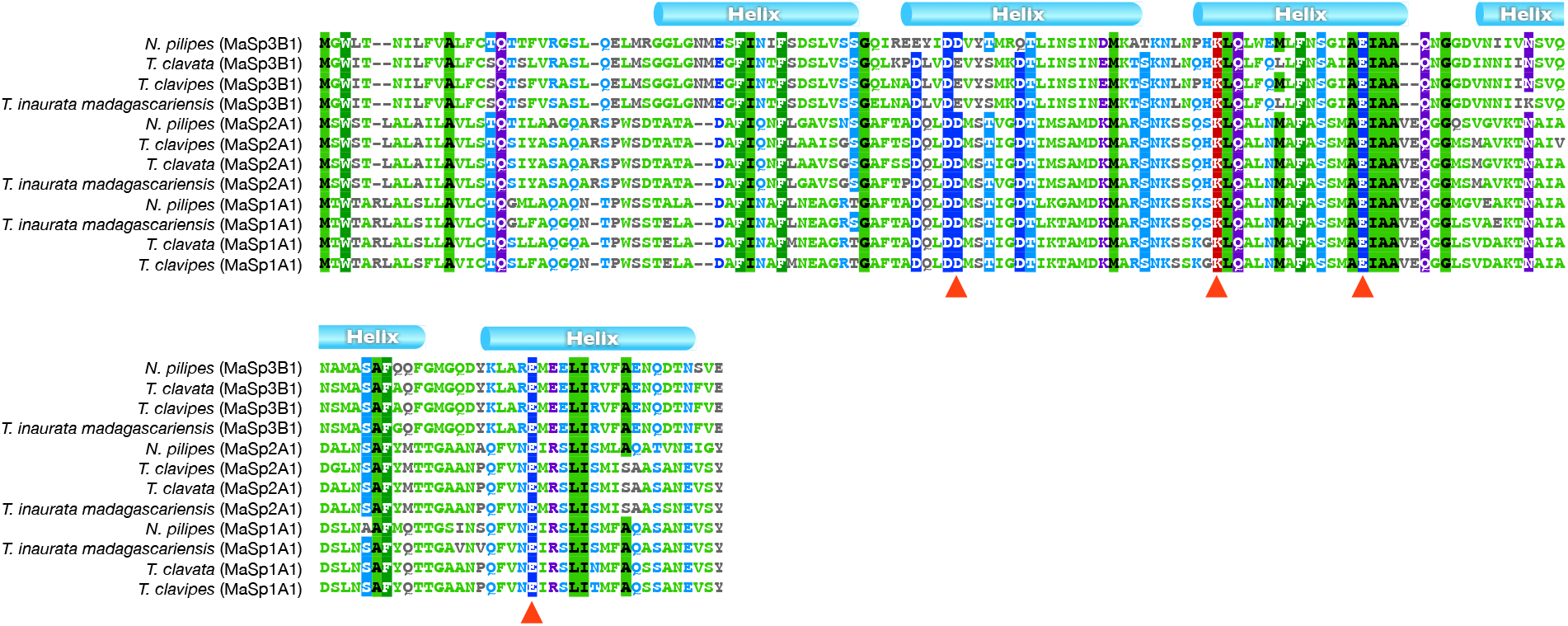
Sequence alignment of MaSp N-terminal regions. This alignment figure confirms which residues in the N-terminal domain (NTD) are shared among MaSp1, 2, and 3. Red triangles indicate the residues that have been suggested to contribute to the NTD-assisted association as D40, K65, E79, and E119 (10–12).

**Table S1.** Raw sequencing data

**Table S2.** Transcriptome data of *Trichonephila clavata*

**Table S3.** Transcriptome data of *Trichonephila clavipes*

**Table S4.** Transcriptome data of *Nephila pilipes*

## Notes

### Competing Interest Statement

H.N., R.O., and D.A.P.M. are employees of Spiber Inc., a venture company selling artificial spider silk products. However, all study design decisions were made by N.K. and K.A. of Keio University, and Spiber Inc. had no role in the study design.

